# Mechanism of RanGTP priming H2A-H2B release from Kap114 in an atypical RanGTP•Kap114•H2A-H2B complex

**DOI:** 10.1101/2022.11.22.517512

**Authors:** Jenny Jiou, Joy M. Shaffer, Natalia E. Bernades, Ho Yee Joyce Fung, Juliana Kikumoto Dias, Sheena D’Arcy, Yuh Min Chook

## Abstract

Previously we showed that the nuclear import receptor Importin-9 wraps around the H2A-H2B core to chaperone and transport it from the cytoplasm to the nucleus. However, unlike most nuclear import systems where RanGTP dissociates cargoes from their importins, RanGTP binds stably to the Importin-9•H2A-H2B complex and formation of the ternary RanGTP•Importin-9•H2A-H2B complex facilitates H2A-H2B release to the assembling nucleosome. It was unclear how RanGTP and the cargo H2A-H2B can bind simultaneously to an importin, and how interactions of the three components position H2A-H2B for nucleosome assembly. Here we show cryo-EM structures of Importin-9•RanGTP and of its yeast homolog Kap114, including Kap114•RanGTP, Kap114•H2A-H2B, and RanGTP•Kap114•H2A-H2B to explain how the conserved Kap114 binds H2A-H2B and RanGTP simultaneously and how the GTPase primes histone transfer to the nucleosome. In the ternary complex, RanGTP binds to the N-terminal repeats of Kap114 in the same manner as in the Kap114/Importin-9•RanGTP complex, and H2A-H2B binds via its acidic patch to the Kap114 C-terminal repeats much like in the Kap114/Importin-9•H2A-H2B complex. Ran binds to a different conformation of Kap114 in the ternary RanGTP•Kap114•H2A-H2B complex. Here, Kap114 no longer contacts the H2A-H2B surface proximal to the H2A docking domain that drives nucleosome assembly, positioning it for transfer to the assembling nucleosome.

**Significance Statement:** Histones and their chaperone networks are typically conserved in eukaryotes. The yeast importin Kap114 and its human homolog Importin-9 share low sequence identity, but both are primary nuclear import receptors for the core histone heterodimer H2A-H2B. Cryo-EM structures of Kap114•H2A-H2B, Kap114•RanGTP and Importin-9•RanGTP complexes show homologous structure and function for Kap114 and Importin-9. In the nucleus, RanGTP binding to Kap114/Imp9•H2A-H2B does not release H2A-H2B, but RanGTP binds to form an atypical ternary complex. Structure of the ternary RanGTP•Kap114•H2A-H2B complex explains how the GTPase and cargo bind simultaneously to Kap114 and how the presence of Ran in the complex primes H2A-H2B transfer to assembling nucleosomes.

## Introduction

In the nucleus, core histones H2A, H2B, H3 and H4 organize DNA into nucleosomes as 147 base pairs of DNA wrapped around the (H3-H4)_2_ tetramer and two H2A-H2B dimers (Luger et al. 1997). Core histones are synthesized in the cytoplasm and folded into heterodimers that are actively transported into the nucleus by nuclear import receptors (importins) of the Karyopherin-β family. Cellular studies in *Saccharomyces cerevisiae* (*Sc*) and proteomic analysis of human cells identified the primary importins for H2A-H2B to be *Sc*Kap114 and its human homolog Importin-9 (Imp9, also known as IPO9) (1, 2). It was previously suggested that negatively-charged importins could act as chaperones for highly positively charged nucleic acid-binding proteins (such as core histones) in addition to transporting them into the nucleus (3). The crystal structure of Imp9 bound to H2A-H2B showed the spiral-shaped HEAT-repeat containing importin wrapping around the globular histone-fold domain of H2A-H2B (4). This mode of interaction occludes large surfaces on H2A-H2B that bind DNA and H3-H4 in the nucleosome (4). Imp9 thus acts as a histone chaperone that shields H2A-H2B from spurious interactions with nucleic acids and proteins while transporting it from the cytoplasm into the nucleus.

After the Kap114/Imp9•H2A-H2B complex is transported into the nucleus across the nuclear pore complex, H2A-H2B still needs to be delivered to assembling nucleosomes. Typically, importin-cargo complexes are dissociated by the GTP-bound Ran GTPase in the nucleus (5–8). High concentration of RanGTP in the nucleus is maintained by Ran guanine nucleotide exchange factor, RCC1, which is chromatin-bound and catalyzes the exchange to RanGTP (9–12). RanGTP often binds importin and releases cargo into the nucleus (13–17). However, nuclear import of H2A-H2B by Imp9 was shown to be an exception to this Ran-mediated cargo release mechanism. RanGTP does not release H2A-H2B from Imp9; instead, RanGTP binds the Imp9•H2A-H2B complex to form a stable RanGTP•Imp9•H2A-H2B complex (4). Within this ternary complex, RanGTP-binding modulates Imp9-H2A-H2B interactions to allow H2A-H2B-DNA interactions and to facilitate deposition of H2A-H2B onto an assembling nucleosome. Pemberton and colleagues also found a stable assembly containing Imp9 homolog Kap114, H2A-H2B, RanGTP and the histone chaperone Nap1 in the yeast nucleus (18).

It was unclear how Kap114/Imp9 binds H2A-H2B and RanGTP simultaneously as a ternary complex of Ran, importin and cargo is atypical among nuclear import complexes. It was also not known how RanGTP-binding changes Kap114/Imp9-H2A-H2B interactions within the ternary complex to facilitate nucleosome assembly. We solved cryo-EM structures of binary complexes Imp9•RanGTP, Kap114•RanGTP and Kap114•H2A-H2B, which collectively show conservation of H2A-H2B and RanGTP recognition by Imp9 and its yeast homolog Kap114. Comparison of the H2A-H2B-*versus* Ran-bound binary complexes shows conformational changes at the N-terminal HEAT repeats of the importins even though there are few overlaps in the importin residues that bind H2A-H2B and RanGTP. The C-terminal HEAT repeats of the importins make extensive interactions with H2A-H2B but make no contact with RanGTP and are thus flexible in the importin•Ran structures. Most importantly, we solved the structure of the RanGTP•Kap114•H2A-H2B ternary complex, which shows that Kap114 binds H2A-H2B differently when RanGTP is also bound to the importin. The Kap114 C-terminal HEAT repeats maintain extensive contacts with the nucleosomal DNA and H3-H4-binding surfaces of H2A-H2B, but the Kap114 N-terminal HEAT repeats no longer contact the remaining nucleosomal DNA-binding surface of H2A-H2B. The exposed H2A-H2B surface allows interactions with DNA, facilitating transfer of H2A-H2B to the assembling nucleosome.

## Results

### Interactions of Kap114 with H2A-H2B and RanGTP

We used pull-down binding assays, electrophoretic mobility shift assays (EMSAs), and analytical ultracentrifugation sedimentation analysis (AUC) to characterize Kap114 binding to H2A-H2B and RanGTP. Consistent with their homologous sequences, architectures and functions, both Kap114 and Imp9 bind to H2A-H2B with sub-nanomolar affinities (Figure S1A-D). Like all importins, they also both bind very tightly to RanGTP (K_D_s < 1nM) (Figure S1E). Pull-down binding assays show that immobilized maltose-binding protein (MBP)-Kap114 fusion bound H2A-H2B and RanGTP simultaneously; RanGTP does not cause displacement of H2A-H2B from Kap114 (Figure 1A, controls in Figure S2A). EMSAs further show that Kap114 bound 1:1 to H2A-H2B or RanGTP and verified formation of a ternary complex of RanGTP•Kap114•H2A-H2B (Figure 1B, Figure S2B). This stable ternary complex sediments at 7.4 S, compared to binary Kap114•RanGTP and Kap114•H2A-H2B complexes that sediment at 6.3 S and 6.7 S respectively (Figure 1C). As a measure of histone chaperone activity of Kap114, we performed DNA competition with Kap114 or Imp9 in the presence and absence of RanGTP (Figure 1D, S2C and D). Both Imp9 and Kap114 bound H2A-H2B and released DNA from DNA•H2A-H2B while Imp9•RanGTP and Kap114•RanGTP did not (Figure 1D, compare lanes 4-6 and 8-10 of Figure S2D). These results are consistent with Padavannil et al. where the presence of RanGTP changes Kap114/Imp9-H2A-H2B chaperoning ability (4). Kap114 and Imp9 are thus similar in having histone chaperone activity, in forming a ternary complex with RanGTP and H2A-H2B, and in RanGTP modulating importin-histone interactions within the ternary complex.

**Figure 1.**
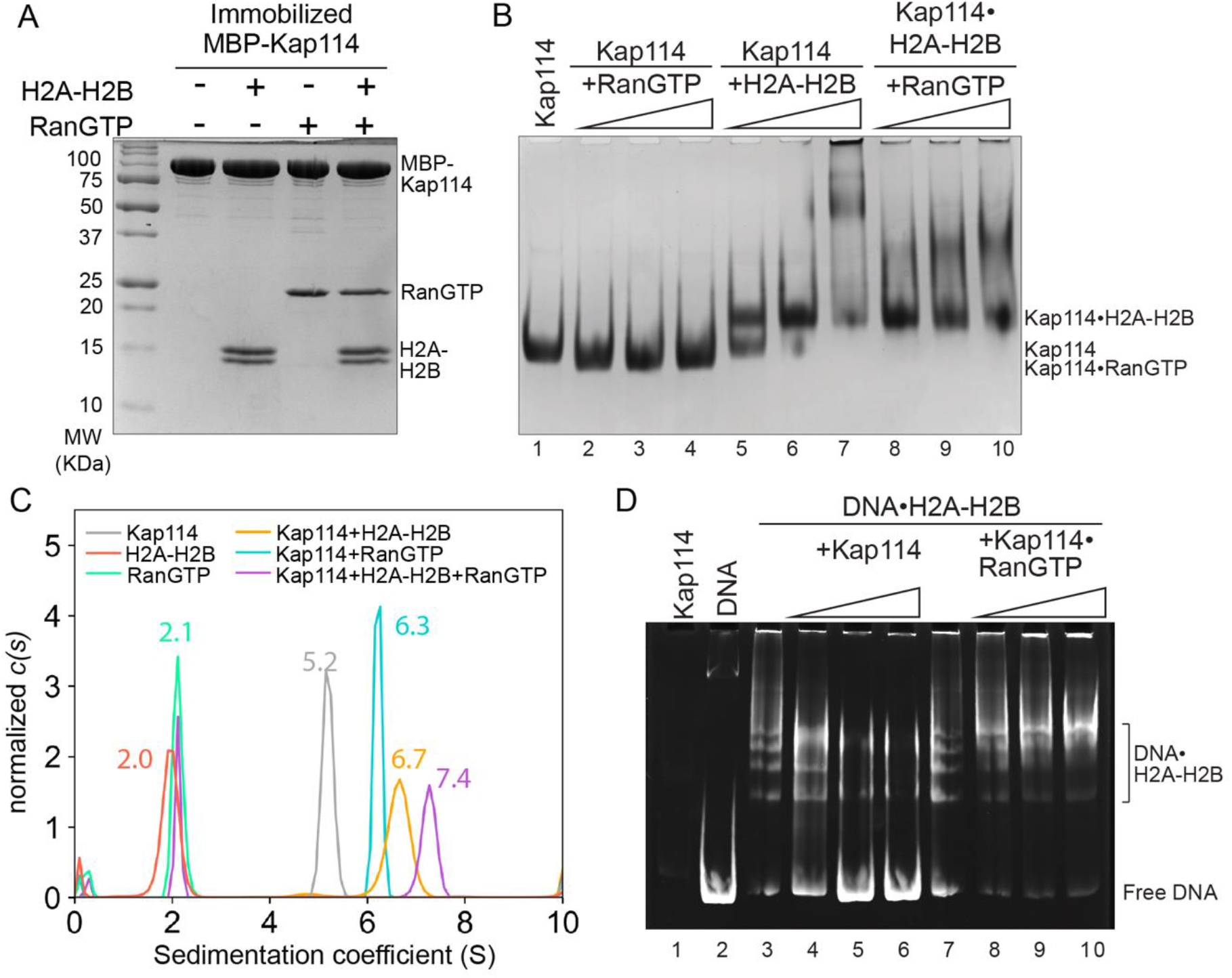
Kap114 binding to H2A-H2B and RanGTP. **(A)** Pull-down binding assay of MBP-Kap114 immobilized on amylose beads with H2A-H2B and RanGTP. After extensive washing, bound proteins were visualized by Coomassie-stained SDS-PAGE. Controls are in Figure S2A. **(B)** Constant Kap114 was titrated with 0.5, 1, or 1.5 molar ratio of RanGTP (lanes 2-4) or H2A-H2B (lane 5-7). Constant Kap114•H2A-H2B was titrated with 0.5, 1, or 1.5 molar ratio of RanGTP (lanes 8-10). Protein was visualized by Coomassie-stained native PAGE. (C) AUC sedimentation profiles of Kap114 (grey), H2A-H2B (red), RanGTP (green), 1:1 molar ratio of Kap114:H2A-H2B (orange), 1:1 molar ratio of Kap114:RanGTP (blue), and 1:1:3 molar ratio of Kap114:H2A-H2B:RanGTP (purple). **(D)** DNA competition assay with Kap114 (lanes 4-6) titrated at 0.5, 1, or 1.5 molar equivalents of H2A-H2B (in a DNA•H2A-H2B 1:7 complex), while Kap114•RanGTP (1:1, lanes 8-10) is titrated at 0.25, 0.5, or 1.0 molar equivalents of H2A-H2B (in a DNA•H2A-H2B 1:7 complex). Native PAGE was visualized with ethidium bromide. Coomassie staining is shown in Figure S2C.

### Structures of binary complexes Imp9•RanGTP, Kap114•RanGTP and Kap114•H2A-H2B

We solved cryo-EM structures of the following binary complexes: Kap114•H2A-H2B (3.2 Å resolution), Kap114•RanGTP (3.5 Å resolution), and Imp9•RanGTP (3.7 Å resolution) (Figure 2A-C, S3 and 4, Table S1 and 2). Imp9 and *Sc*Kap114 are 21.2% identical in sequence and share all key structural features, including 20 HEAT repeats (h1-h20, each containing antiparallel helices a and b) and three long loops (the h8 loop, the h18-h19 loop, and the acidic h19 loop) (Figure S5). Structures of Imp9 and Kap114 bound to H2A-H2B or to RanGTP are very similar.

**Figure 2.**
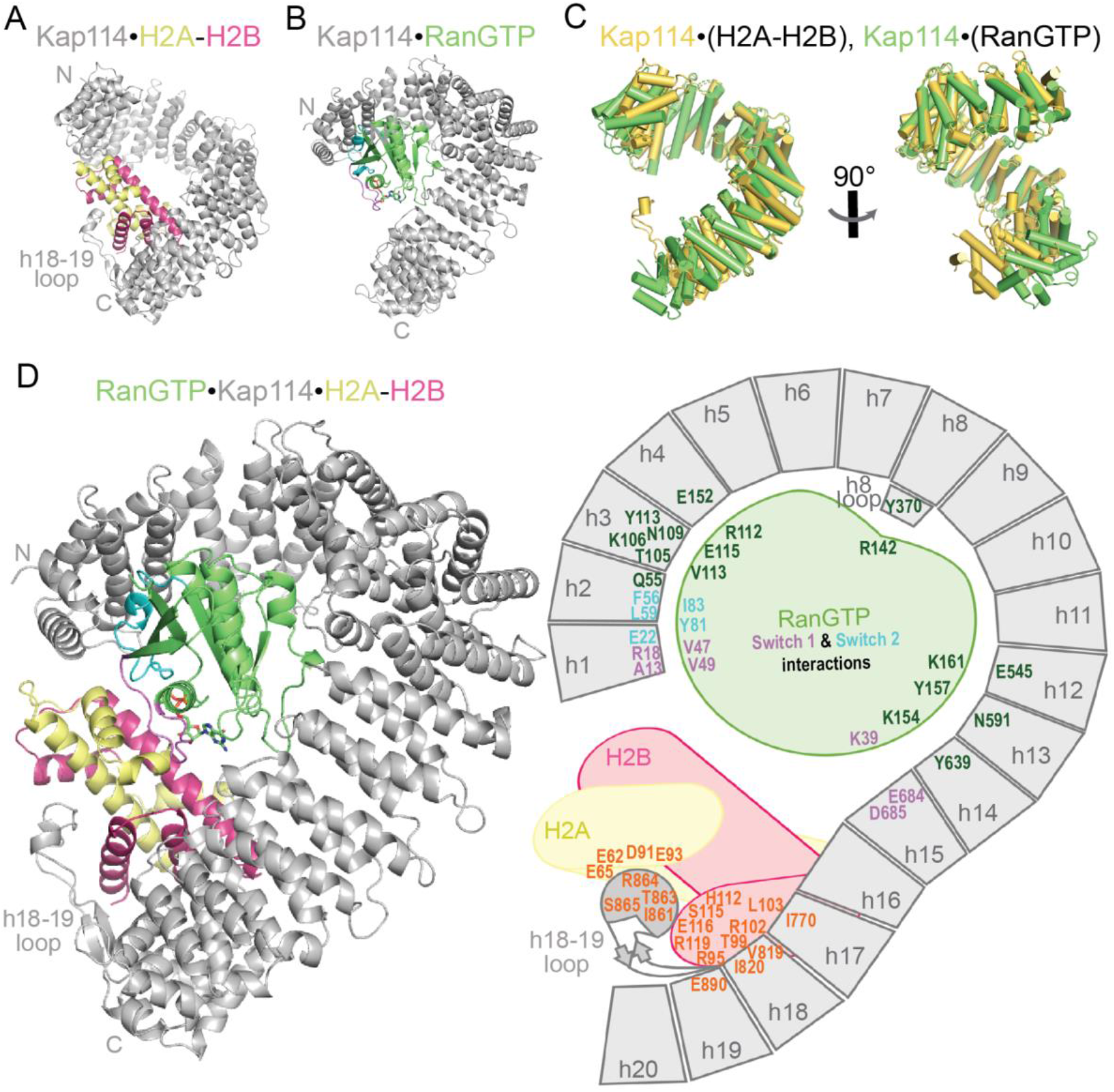
Cryo-EM structures of Kap114•H2A-H2B, Kap114•RanGTP and RanGTP•Kap114•H2A-H2B. (**A**) Cartoon representation of Kap114(gray)•H2A(yellow)-H2B(red). (**B**) Cartoon representation of Kap114•RanGTP. Switch 1 (violet), Switch 2 (cyan) and GTP (sticks colored by atom) of RanGTP (green) are shown. (**C**) Superposition of Kap114 from Kap114•H2A-H2B (yellow) and Kap114•RanGTP (green) aligned at HEAT repeats h5-h13 (RMSD=0.515 Å). (**D**) Cartoon representation of RanGTP•Kap114•H2A-H2B on the left and a schematic with interacting residues indicated on the right. H2A-H2B interactions are in orange and RanGTP interactions are in violet (Switch 1), cyan (Switch 2), and green.

When bound to H2A-H2B, Kap114 adopts a compact, super-helical architecture that is similar to Imp9 bound to H2A-H2B (Figure 2A, Figure S6). The Kap114 superhelix wraps around the H2A-H2B core using interactions conserved with Imp9 (Figure S6–7). The C-terminal ends of h3-h5 b-helices, the loops that follow these helices and helix h2b, the loops of h17 and h18 repeats, the C-terminal end of h19a helix, and the long h18-h19 loop, all contact H2A-H2B. Sequence comparison of Kap114 and Imp9 reveals greatest identity in the C-terminal HEAT repeats (27.3% identity aligning 319 residues of h16-h20); the h18-h19 loop is especially conserved. The N-terminal HEAT repeats (20.8% identity aligning 240 residues of h1-h5) and the central HEAT repeats (17.9% identity aligning 549 residues of h6-h15) are also conserved (Figure S5). This sequence conservation is consistent with similar H2A-H2B binding by Kap114 and Imp9 at both their C- and N-terminal HEAT repeats (detail interactions in Figure S6).

RanGTP-bound structures of Kap114 and Imp9 are also very similar and are like the many previously reported importin•Ran structures (Figure 2B, S8 and 9) (14, 19–25). Kap114 and Imp9 primarily use the b-helices of h1-h4 to bind Switch 1, Switch 2 and helix α3 of RanGTP. Other interactions involve the extended h8 loops and h12-15 of the importins contacting helix α4 and Switch 1 of RanGTP, respectively. The C-terminal repeats of Kap114 (h16-h20) make no contact with RanGTP, consistent with three equally-represented classes of Kap114•RanGTP cryo-EM particles that differ in the orientations of their C-terminal repeats (Figure S4; only class 1 structure is determined and shown in Figure 2B). The flexible C-terminal repeats of Kap114 are also consistent with the lack of density for the likely very flexible h16-h20 of Imp9•RanGTP (Figure S3B).

Comparison of the Kap114•H2A-H2B and Kap114•RanGTP structures show minimal overlap between residues that contact H2A-H2B *versus* RanGTP (Figure S5). However, the importin conformations, especially at the N-terminal regions, are quite different when bound to the two ligands (Figure 2A-C). The RanGTP-bound Kap114 superhelix is wider and has a shorter pitch than the H2A-H2B-bound one (Figure 2C). Alignments of HEAT repeats between the Kap114•H2A-H2B and Kap114•RanGTP structures reveal rigid body regions and hinges between them that describe the conformational differences (Figure S10A). Repeats h5-h13 are similar in the two structures (RMSD=0.515 Å for 378 of 430 Cα atoms aligned), suggesting that this is a rigid body block. Meanwhile, high RMSDs when aligning consecutive repeats suggest that h4-h5, h13-h14 and h16-h17 are hinges about which groups of HEAT repeats rotate. These groups of repeats are 1) h1-h4 that contact H2A-H2B and RanGTP, 2) the invariant h5-h13 core and 3) the h17-h20 repeats that make no contact with RanGTP and are very flexible in Kap114•RanGTP but make extensive contacts with H2A-H2B. The hinges at h4-h5 facilitate movement of the N-terminal repeats (h1-h4) relative to the central core repeats (h5-h13) while the h13-h14 and h16-h17 hinges facilitate movement of the C-terminal repeats (h17-h20) relative to the central core repeats (Figure 2C, S10A). Similar features of conformational change are observed for Imp9•H2A-H2B and Imp9•RanGTP (Figure S10B).

### Structure of the RanGTP•Kap114•H2A-H2B ternary complex

It was unclear from the binary structures how the ternary complex is arranged when Kap114 binds RanGTP and H2A-H2B simultaneously. We assembled the RanGTP•Kap114•H2A-H2B complex for structure determination by cryo-EM. The data collected produced three classes of ternary complexes that differ in the orientations of Kap114 repeats h16-h20. We solved the structure of the most highly populated class 3 (Figure 2D, S11 and 12). The structure of this most open/fanned-out conformation of the RanGTP•Kap114•H2A-H2B complex was determined to 3.3 Å resolution (Figure 2D and S12, Table S1).

The RanGTP•Kap114•H2A-H2B structure (Figure 2D) shows the N-terminal half of Kap114 superhelix (composed of h1-h15) binding RanGTP by adopting the same conformation (RMSD=0.625 Å, 587 of 701 Cα atoms aligned) and making the same interactions with the GTPase as the binary Kap114•RanGTP structure (Figure 3A, B and S9). Residues in the Kap114 h1-h4 repeats that contacted H2A-H2B in Kap114•H2A-H2B are still accessible in the ternary complex but this region is now further away from the bound histone, with minimally ~5 Å between H2A-H2B and the h2-3loop of Kap114 (Figure 3C). There is a drastic decrease in interaction at the N-terminal repeats of Kap114 when compared to Kap114•H2A-H2B (Figure 3C, D). Only few contacts are present between the RanGTP switches with H2A but the lack of H2A-H2B side chain densities, consistent with local resolution of 4.5 Å for H2A-H2B as shown in Figure S12A-C, suggests that these Ran-histone interactions in the ternary complex are transient (Figure 3C). This newly exposed surface of H2A-H2B is also proximal to the H2A C-terminal tail that docks onto H3-H4 in the nucleosome, although this tail is not visible in either structure.

**Figure 3.**
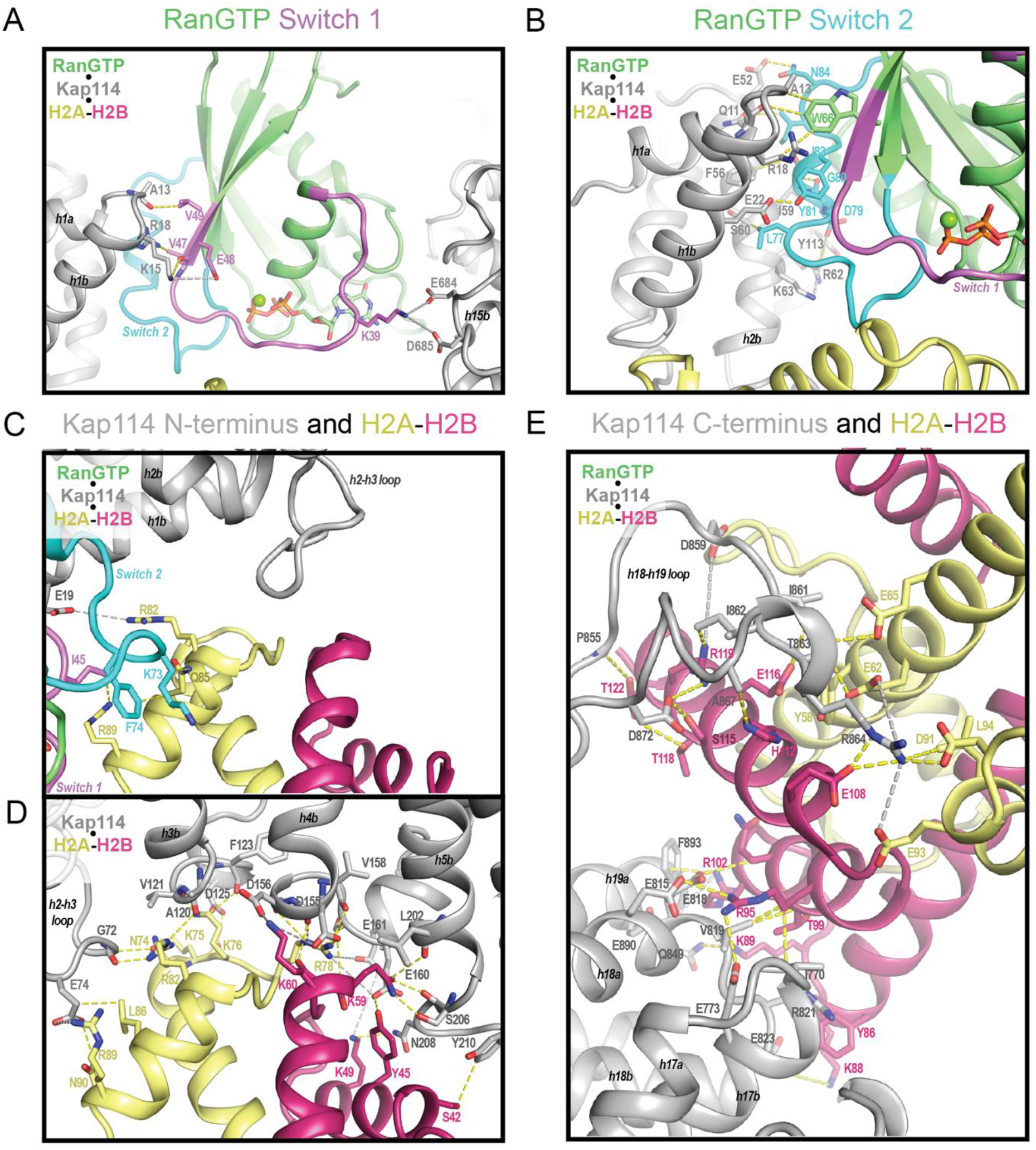
Interactions of the ternary RanGTP•Kap114•H2A-H2B complex. (**A**) Detail interactions of Kap114 with RanGTP Switch 1, (**B**) Switch 2, and (**C**) RanGTP, H2A-H2B and N-terminal repeats and of Kap114. (**D**) In same orientation as C, Kap114 interactions with H2A-H2B in the binary structure. (**E**) The C-terminal repeats of Kap114 in the binary Kap114•H2A-H2B complex. In all panels, Kap114 is gray, RanGTP is green with Switch 1 in violet and Switch 2 in cyan, H2A is yellow, and H2B is red. Dashed lines represent interactions <4 Å (yellow) and long-range electrostatic interactions <8 Å (gray).

The region C-terminal of h16 in Kap114 does not contact RanGTP but adopts the same conformation and maintains the same extensive contacts with H2A-H2B as in the Kap114/Imp9•H2A-H2B structures (Figure 3E and S6). The C-terminal region of Kap114, with H2A-H2B bound, adopts different orientations relative to the N-terminal half of the importin in the different cryo-EM classes of particles (Figure 2D and S11). The movements of the histone-bound Kap114 C-terminal region likely occur about the hinge at h16-h17.

### The GTPase in RanGTP•Kap114•H2A-H2B primes transfer of H2A-H2B to nucleosome

We performed nucleosome assembly assays where we titrated H2A-H2B and Kap114 in the presence or absence of RanGTP into tetrasomes (Figure 4A and S13). Like Imp9, Kap114 facilitated H2A-H2B deposition, forming a nucleosome, only in the presence of RanGTP (Figure 4A and S13). Without RanGTP, Kap114/Imp9 inhibited H2A-H2B deposition. The RanGTP-induced release of the contacts between H2A-H2B and the N-terminal HEAT repeats of Kap114/Imp9 clearly makes H2A-H2B more available to form nucleosomal interactions with DNA and H3-H4. These results are consistent with our structural data.

**Figure 4.**
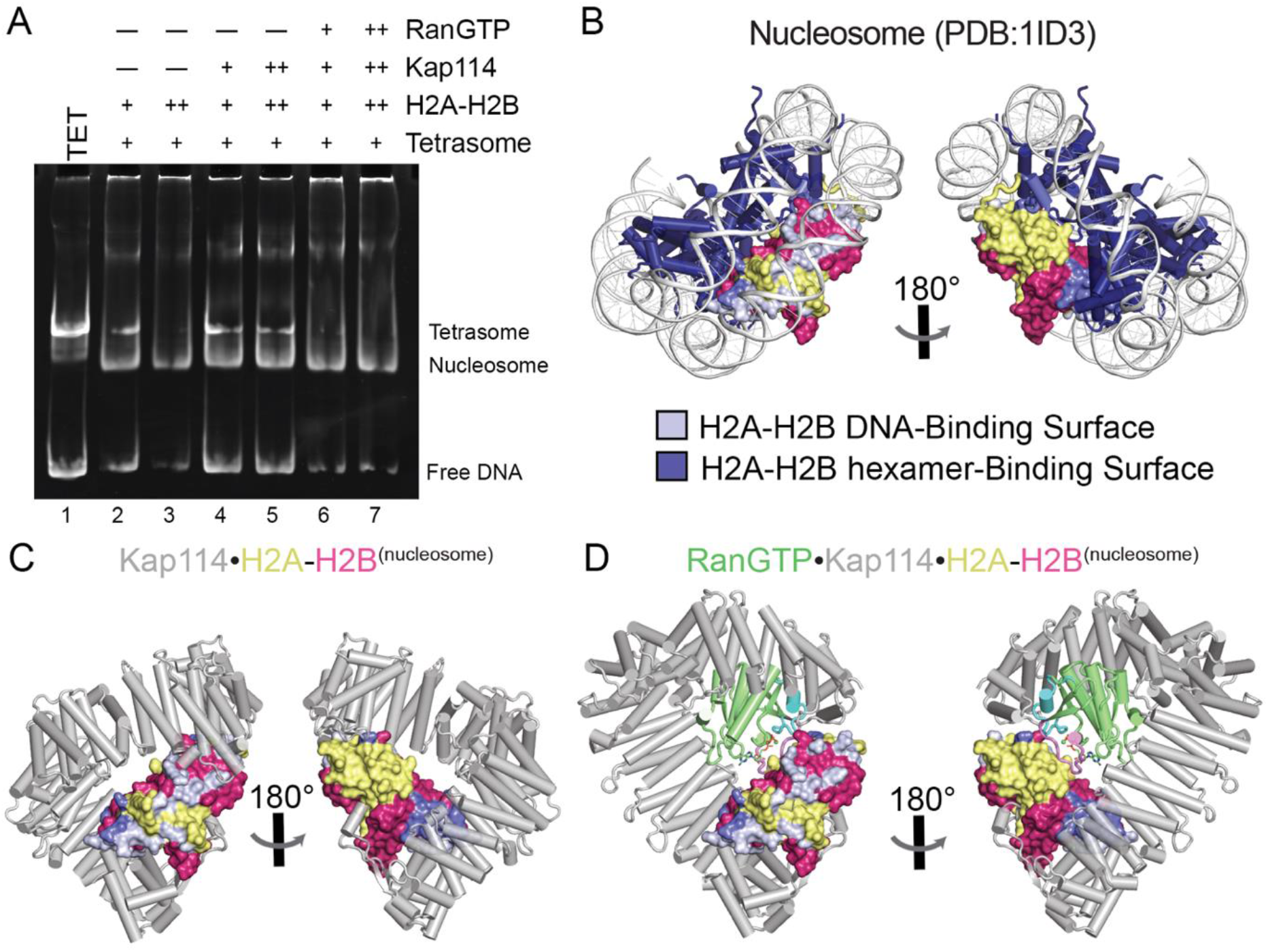
Nucleosome assembly assay and surface representation of nucleosomal binding surfaces on H2A-H2B. **(A)** Nucleosome assembly assay where either H2A-H2B, Kap114•H2A-H2B or RanGTP•Kap114•H2A-H2B is titrated in molar equivalents of 2.0 and 3.0 to tetrasome (TET; 1.25 μM). Ethidium bromide-stained gel is shown. The same gel is Coomassie-stained in Figure S13A. **(B)** Nucleosome (PDB 1ID3) highlighting a single copy of H2A-H2B (yellow and red, respectively) and its interactions. One H2A-H2B core is shown as surface while the H2A C-terminal tail is shown as cartoon (as it is not visible in the importin structures). The DNA is white with its interactions on H2A-H2B colored blue-white. The histone hexamer, H2A-H2B+(H3-H4)_2_, is navy with its interaction on H2A-H2B colored slate-blue. The surface of the nucleosome H2A-H2B core and its interaction colors are aligned and docked into (**C**) Kap114•H2A-H2B and (**D**) RanGTP•Kap114•H2A-H2B. Left and right views are 180° rotations.

We compared the location of nucleosomal interfaces of H2A-H2B to Kap114•H2A-H2B and RanGTP•Kap114•H2A-H2B structures to understand how RanGTP facilitates H2A-H2B release to the assembling nucleosome (Figure 4B-D). In Kap114•H2A-H2B, a large portion of the nucleosomal DNA-binding surface of H2A-H2B is obstructed by the N- and C-terminal regions of Kap114 (Figure 4B and C). The region of H2A-H2B that interfaces with H3-H4 through a four-helix bundle in the nucleosomes is also covered by the C-terminal region of Kap114 (Figure 4B and C). This extensive occlusion in the binary Kap114/Imp9•H2A-H2B complex explains the poor ability to release H2A-H2B to DNA (Figure 1D) or tetrasomes (Figure 4A, S13A and B). Occlusion of H2A-H2B is altered in the ternary RanGTP•Kap114•H2A-H2B structure. Although the Kap114 C-terminal repeats still cover part of the nucleosomal DNA and H3-H4-binding surfaces of H2A-H2B, the Kap114 N-terminal repeats no longer engage the remaining nucleosomal DNA-binding surface of H2A-H2B. This exposure of a long stretch of DNA-binding residues proximal to the H2A C-terminal tail, makes H2A-H2B more available for potential capture by DNA (Figure 1D) or the assembling nucleosome (Figure 4B and D). These structural observations are consistent with the nucleosome assembly assays where the RanGTP•Kap114/Imp9•H2A-H2B complex effectively deposits H2A-H2B onto assembling nucleosomes (Figure 4A and S12).

## Discussion

Kap114 behaves like Imp9 in all our structural, biophysical and biochemical analyses. Their structures bound to H2A-H2B or RanGTP are very similar with only minor differences, such as no interaction between Kap114 and the H2B N-terminal tail compared to 3-5 tail residues seen bound to Imp9 (4). The lack of H2B tail contacts with Kap114 is consistent with previous findings that the histone tails are not important for Imp9 binding, and that the removal of the histone tails does not affect their nuclear import (4, 26). Both Imp9 and Kap114 occlude the nucleosomal DNA-binding regions of H2A-H2B with their N-terminal HEAT repeats, and the nucleosomal DNA- and histone-binding sites with their C-terminal HEAT repeats. Such extensive interfaces render Imp9 and Kap114 effective H2A-H2B chaperones that compete interactions between H2A-H2B and DNA.

However, the very high affinity and extensive importin-histone interactions could make it difficult to release H2A-H2B in the nucleus. Indeed, unlike other import cargos that are easily dissociated from their importins by RanGTP, the GTPase cannot release H2A-H2B from Kap114 or Imp9 except in the presence of assembling nucleosomes. The stable RanGTP•Kap114•H2A-H2B complex allows Kap114 to continue to chaperone H2A-H2B in the nucleus, consistent with the generally accepted notion that there is no free H2A-H2B in the cell (27, 28). H2A-H2B release becomes targeted as conformational changes in RanGTP•Kap114•H2A-H2B release contacts between H2A-H2B and the N-terminal HEAT repeats of Kap114 to make H2A-H2B available for nucleosomal interactions with DNA and H3-H4. These findings are validated and extended to Imp9 by solution HDX analysis (manuscript co-submitted, Shaffer et al.).

Although we have provided a structural understanding for how RanGTP primes the release of H2A-H2B from Kap114/Imp9 to assembling nucleosomes, many questions remain about this process in cells. First, it is unclear if additional factors are involved. Second, sumoylation of Kap114 was reported to improve the ability of RanGTP to release cargoes Sua7 (also known as TFIIb) and TBP (29). The sumoylation site, mapped to residue K909, is in the long disordered acidic h19 loop that is not modeled in any of our Kap114 structures. Deletion of the h19 loop in Imp9 did not significantly affect H2A-H2B binding (4); it remains unclear how sumoylation regulates RanGTP-mediated release of H2A-H2B from Kap114/Imp9. Third, we showed that RanGTP•Kap114/Imp9 cannot disassemble H2A-H2B from nucleosomes but it is not known if nuclear Kap114/Imp9 in the presence of RanGTP may chaperone H2A-H2B after it is removed from the nucleosome by remodelers during replication, transcription or DNA repair (30, 31). Furthermore, the importin-histone interface in RanGTP•Kap114/Imp9•H2A-H2B involves the H2A-H2B acidic patch, which also mediates inter-nucleosome contacts that are important for higher-order chromatin structures (32, 33). Kap114/Imp9 may have potential roles in modulating higher-order chromatin structure and function. Finally, Kap114 associates with Nap1 and H2A-H2B in both the cytoplasm and nucleus (1, 18). Imp9 also co-purified with Nap1L1 and Nap1L4 in mammalian cells (34–37). The Nap1-binding site of H2A-H2B overlaps with the N-terminal Kap114-binding site but not the C-terminal binding site (4, 38), which may be how Kap114 binds Nap1-bound H2A-H2B. It is unclear whether H2A-H2B is transferred between Nap1 and Kap114 akin to transfers in the histone chaperone network (39–42) or H2A-H2B is co-chaperoned by Nap1 and Kap114 like in ASF1•H3-H4•Importin-4 (25).

We have revealed how Kap114 chaperones H2A-H2B in the cytoplasm and in the nucleus; the latter in the form of the RanGTP•Kap114•H2A-H2B complex. Our previous study of Imp9 binding to H2A-H2B and solution HDX analysis of Imp9 show that this importin uses the same mechanism (4, manuscript co-submitted, Shaffer et al.). This knowledge is the foundation to understand how additional molecular players contribute to H2A-H2B cytoplasmic processing and deposition into the nucleosome.

## Methods

### Constructs, Protein Expression and Purification

*Sc*Kap114 was cloned into two modified vectors: pGEX-4T3 (GE Healthcare) and pmalE (New England BioLabs). The pGEX-43T was modified to have a TEV cleavage site inserted between the GST tag and Kap114. The pmalE was modified to have a His-tag at the N-terminus of MBP and a TEV cleavage site after the MBP.

The plasmid was transformed into BL21-Gold(DE3) and plated on Luria Broth (LB) agar with ampicillin for selection. Kap114 was expressed using 6 L of LB media and induced with 0.5 mM isopropyl β-D-1-thiogalactopyranoside (IPTG) for 16 h at 20°C. Cells were spun at 4,000 rpm with a Sorvall BP8 (Thermo Fisher) for 30 min at 4°C and resuspended in lysis buffer (20 mM Tris-HCl pH 7.5, 1 M NaCl, 15% (v/v) glycerol, 2 mM dithiothreitol (DTT for GST-Kap114) or 2 mM β-mercaptoethanol (BME for His_6_MBP-Kap114), cOmplete EDTA-free Protease Inhibitor Cocktail (Roche Applied Science). Cells were lysed with an Emulsiflex-C5 cell homogenizer (Avestin) and centrifuged in an Avanti J-26 XPI with a JA-25.50 rotor (Beckman Coulter) at 20,000 RCF for 1 h at 4°C. The supernatant was decanted into a gravity, affinity column. GST-Kap114 was purified with Glutathione Sepharose 4B (Cytiva) and the GST tag was cleaved on column using TEV protease. His_6_MBP-Kap114 was purified with Ni^2+^-NTA Agarose (Qiagen). Once eluted, the Kap114 or His_6_MBP-Kap114 was further purified by ion-exchange chromatography using HiTrap Q HP (Cytiva), and gel filtration chromatography using Superdex 200 16/600 (Cytiva). Proteins were stored in the gel filtration buffer containing 20 mM Tris-HCl pH 7.5, 150 mM NaCl, 2 mM DTT.

Human Imp9 and *Sc*Ran (Gsp1 residues 1-179 with Q71L) were expressed and purified as previously described (4). *SclXenopus laevis (Xl*) histones were obtained from The Histone Source and refolded according to established protocol (32). 147 bp DNA was purified and assembled into tetrasomes using established protocols (32).

mNeonGreen was sub-cloned from pL0M-S-mNeonGreen-EC18153 into pET28a along with GSP1(1-179, Q71L). pL0M-S-mNeonGreen-EC18153 was a gift from Julian Hibberd (RRID:Addgene 137075). mNeonGreen-GSP1 was expressed in Rosetta(DE3)pLysS cells using 2 L of LB media with kanamycin and chloramphenicol for selection. The cells were induced with 0.25 mM IPTG for 12 h at 18°C. Protein was purified as for GSP1. H2A-K119C was labeled with XFD488 (ATT Bioquest) and prepared according to the manufacturer’s protocol.

### Pull-down Binding Assays

Pull-down binding assays were performed by immobilizing purified His_6_MBP or His_6_MBP-Kap114 on amylose resin (New England BioLabs). Resin was stored in Binding Assay (BA) buffer containing 20 mM Tris-HCl pH 7.5, 150 mM NaCl, 10% (v/v) glycerol and 2 mM DTT forming a 50% amylose resin BA slurry. 100 μL of slurry with a total solution volume of 50 μL BA buffer was brought up to a total solution volume of 100 μL with a final concentration of 10 μM His_6_MBP or His_6_MBP-Kap114, 50 μM *Sc*H2A-H2B, and/or 50 μM RanGTP. RanGTP was added after a 30 min pre-incubation period of the other components and equilibrated for another 30 min at room temperature. Amylose resin was pelleted at 16,000 RCF for 1 min at 4°C using an Eppendorf Centrifuge 5415 R and washed 3 times with 600 μL of BA buffer stored at 4°C with excess solution carefully aspirated. 100 μL of 2x Laemmli sample buffer was added, and BA samples were boiled for 5 min. 10 μL of sample was loaded onto a 12% SDS-PAGE gel. Gels were visualized using Coomassie stain.

### Electrophoretic Mobility Shift Assays

For Electromobility Shift Assays (EMSAs) with Imp9, one component was held constant at 10 μM while the other was titrated. For EMSAs with Kap114, one component was held constant at 5 μM, while the other was titrated. Proteins were dialyzed into the same buffer overnight (20 mM HEPES, pH 7.5, 150 mM NaCl, 2 mM MgAcetate, 2 mM Tris(2-carboxyethyl)phosphine hydrochloride (TCEP), 10% (v/v) glycerol). Samples were separated by 5% native PAGE. Gels were run for 100 min at 150 V at 4 °C in 0.5x TBE (40 mM Tris-HCl pH 8.4, 45 mM boric acid, 1 mM Ethylenediaminetetraacetic acid (EDTA)). Gels were stained with Coomassie. Gel shown was one of ≥ 3 repeats.

### Analytical Ultracentrifugation

Sedimentation velocity experiments were performed as described in Padavannil et al. (4). Individual proteins were dialyzed into Analytical Ultracentrifugation (AUC) buffer containing 20 mM Tris-HCl pH 7.5, 150 mM NaCl, 2 mM MgCl_2_ and 2 mM TCEP. 500 μL of AUC sample was equilibrated overnight at 4°C. The AUC sample chamber contained 400 μL of the following: 1) 3 μM Kap114, 2) 10 μM ScH2A-H2B, 3) 10 μM RanGTP, 4) 3 μM Kap114 + 3 μM H2A-H2B, 5) 3 μM Kap114 + 3 μM RanGTP, 6) 3 μM Kap114 + 3 μM H2A-H2B + 10 μM RanGTP. Sedimentation coefficients were measured by monitoring absorbance at 280 nm in a Beckman-Coulter Optima XL-1 Analytical Ultracentrifuge. The Beckman data time stamps were corrected using REDATE (43). SEDNTERP was used to calculate the buffer density, buffer viscosity, and protein partial-specific volumes (44). SEDFIT was used to calculate sedimentation coefficient distributions c(S) where the regularization calculated a confidence level of 0.68 was used, time-independent noise elements were accounted for, and at a resolution of 100 (45). SEDFIT was also used to obtain the sedimentation coefficient by integration of c(S) and the frictional ratios by refining the fit of the model. The data were plotted using GUSSI (46).

### DNA Competition Assays

Imp9, Kap114, Imp9•RanGTP, or Kap114•RanGTP were titrated at various molar equivalents of *Xl*H2A-H2B where H2A-H2B was in a 7:1 complex with 147 bp Widom 601 DNA (10.5 μM H2A-H2B and 1.5 μM DNA). Binary complexes were equimolar Imp9 and RanGTP or Kap114 and RanGTP added together without further purification. Proteins were dialyzed in the same buffer overnight (20 mM HEPES, pH 7.5, 300 mM NaCl, 2 mM MgAcetate, 2 mM TCEP, 10% (v/v) glycerol). Samples were separated by 5% native PAGE. Gels were run for 75 min at 150 V at 4 °C in 0.5x TBE. Gels were stained with ethidium bromide and then Coomassie. Gel shown was one of ≥ 3 repeats.

### Fluorescent Polarization

Fluorescence polarization (FP) assays were performed in a 384-well format as previously described (47). Kap114, Imp9 were dialyzed overnight into 20 mM HEPES pH 7.5, 150 mM NaCl, 2 mM MgCl_2_, 2 mM TCEP, 10% (v/v) glycerol. Kap114 or Imp9 were serially diluted with buffer and mixed with ^XFD488^H2A-H2B or mNeonGreen-RanGTP at indicated concentrations. Triplicate reactions were analyzed in black-bottom plates (Corning) and data were collected in a CLARIOstar plus plate reader (BMG Labtech) equipped with dichroic filter LP 504. Measurement was performed with top optics with an excitation range of 482-16 nm and emission range of 530-40 nm, 50 flashes per well with a 0.1 s settling time. Gain and focal height was adjusted using well with ^XFD488^H2A-H2B and the lowest concentration of Kap114 to target mP of 200 and kept constant for the rest of the measurements unless otherwise noted. Data were analyzed in PALMIST (48) and fitted with a 1:1 binding model, using error surface projection method to calculate the 95% confidence intervals of the fitted data. Fitted data were exported and plotted in GUSSI.

### Cryo-EM sample preparation

For Kap114•H2A-H2B and Kap114•RanGTP, individual proteins were buffer exchanged into cryo-EM buffer containing 20 mM Tris-HCl pH 7.5, 300 mM NaCl, 2 mM MgCl_2_ and 1 mM TCEP and flash frozen at 8 mg/mL. Kap114•H2A-H2B and Kap114•RanGTP were made by mixing Kap114 with *Sc*H2A-H2B or RanGTP, resulting in a 1 Kap114 to 1.2 H2A-H2B or RanGTP molar ratio at 8 mg/mL. Kap114 complexes were diluted two-fold with cryo-EM buffer plus NP-40 at a final concentration of 0.1% (w/v). Samples of Imp9•RanGTP were buffer exchanged into 50 mM Tris-HCl pH 7.5, 150 mM NaCl and 0.1% (w/v) NP-40 at a final protein concentration of 3 mg/ml. 4 μL of Kap114•H2A-H2B/RanGTP or 3.5 μL of Imp9•RanGTP was applied to a 300 mesh copper grid (Quantifoil R1.2/1.3) that was glow-discharged using a PELCO easiGlow glow discharge apparatus for 30 mA/30 s on top of a metal grid holder (Ted Pella). Excess sample was blotted 3 s before plunge-freezing in a ThermoFisher Vitrobot System at 4°C with 95% humidity.

RanGTP•Kap114•H2A-H2B was crosslinked on a Superdex 200 10/300 Increase column equilibrated with cryo-EM buffer. We injected 500 μL of 0.05% (w/v) glutaraldehyde with 3 mL of buffer followed by 500 μL of an equimolar mix of RanGTP, Kap114, and H2A-H2B. We collected 0.5 mL fractions. The ternary complex was concentrated to 8.6 mg/mL and was flash frozen in liquid nitrogen. The complex was diluted two-fold with cryoEM buffer plus Tween-20 (final concentration of 0.00125% (w/v)). The grid was glow-discharged, blotted, and plunge-frozen in a similar manner as the binary complexes.

### Cryo-EM Data Collection

Cryo-EM data for Kap114•H2A-H2B and Kap114•RanGTP were collected at the Pacific Northwest Cryo-EM Center on a Titan Krios at 300 kV with a Gatan K3 detector in correlated double sampling super-resolution mode at a magnification of 81,000x corresponding to a pixel size of 0.5295 Å. Each movie was recorded for a total of 58 frames over 3.475 s with an exposure rate of 15 electrons/pixel/s (total dose of 50 e^-^/Å^2^). Datasets were collected using SerialEM (49) software with a defocus range of −0.8 and −2.5 μm.

Cryo-EM data collection for Imp9•RanGTP and RanGTP•Kap114•H2A-H2B were performed at the UT Southwestern Cryo-Electron Microscopy Facility on a Titan Krios at 300 kV with a Gatan K3 detector in correlated double sampling super-resolution mode at a magnification of 105,000x corresponding to a pixel size of 0.415 Å. Each movie was recorded for a total of 60 frames over 5.4 s with an exposure rate of 7.8 electrons/pixel/s (total dose of 52 e^-^/Å^2^). The datasets were collected using SerialEM with a defocus range of −0.9 and −2.4 μm.

### Cryo-EM Data Processing

A total of 9,158 movies were collected for Kap114•H2A-H2B and 5,744 movies were collected for Kap114•RanGTP. Both datasets were processed using cryoSPARC (50) where they were first subjected to Patch Motion Correction (Kap114•H2A-H2B unbinned and Kap114•RanGTP binned twice) and Patch CTF Estimation. The Blob Picker was implemented on 50 micrographs to pick all possible particles with little bias and this small set of particles were subjected to 2D Classification to generate 2D templates where a subset of templates was used in Template Picker. 7,006,373 particles were picked from the Kap114•H2A-H2B dataset, and 7,640,457 particles were picked from the Kap114•RanGTP dataset. 671,257 particles of Kap114•H2A-H2B and 953,500 particles of Kap114•RanGTP were selected after five rounds of 2D Classification. The Kap114•H2A-H2B particles were then sorted into four 3D classes while the Kap114•RanGTP particles were sorted into six 3D classes using Ab-initio reconstruction followed by Heterogeneous Refinement. The particles from two Kap114•H2A-H2B 3D classes were combined to give 554,529 particles for Non-uniform Refinement which yielded a 3.21 Å resolution map. Three classes of Kap114•RanGTP - Class 1 with 277,305 particles, Class 2 with 258,006 particles and Class 3 with 259,220 particles - were individually subjected to Non-uniform Refinement. We analyzed the local resolution at a 0.5 FSC threshold and determined that Class 1 was the best for model building.

4,767 movies were collected for the RanGTP•Kap114•H2A-H2B complex. This dataset was also processed using cryoSPARC. The movies were subjected to Patch Motion Correction (binned twice) and Patch CTF Estimation. Generation of templates was performed as for the binary complexes and 3,401,608 particles were extracted from the Template Picks. After five rounds of 2D Classification, 604,835 particles were further processed into five Ab-Initio classes and five Heterogeneous Refinement classes. Two of the five classes contained the complex of interest and were subjected to another round of Ab-Initio and Heterogeneous Refinement which generated four new classes. From there, three classes were chosen Non-uniform Refinement: Class 1 with 3.50 Å resolution using 114,232 particles, Class 2 with 3.49 Å resolution using 95,715 particles, and Class 3 with 3.28 Å resolution using 134,915 particles. All maps had local resolution analyzed at a 0.5 FSC threshold, and Class 3 was the best one for model building.

4,920 movies were collected for Imp9•RanGTP and processed using cryoSPARC. A ½ F-crop factor was applied during motion correction followed by patch CTF estimation. A small set of ~20 frames was used to generate the initial template for particle picking. 2,474,170 particles were initially extracted from all the micrographs. The first round of 2D classification produced 904,566 particles that were subjected to another three rounds of 2D classification. 232,798 particles were included for Ab-Initio modeling followed by heterogeneous refinement. Non-uniform refinement was carried out to generate the final 3.7 Å resolution map.

### Cryo-EM Model Building, Refinement, and Analysis

The Kap114 proteins in both Kap114•H2A-H2B and Kap114•RanGTP structures were built using coordinates of unliganded Kap114 from the crystal structure PDB 6AHO (51). The *Sc*H2A-H2B in Kap114•H2A-H2B was built using *Xl*H2A-H2B coordinates from the Imp9•H2A-H2B crystal structure PDB 6N1Z (4) and the RanGTP in Kap114•RanGTP built using coordinates of canine Ran in Kap121•RanGTP crystal structure PDB 3W3Z (22). The Imp9•RanGTP structure was also built using coordinates from PDB 6N1Z and PDB 3W3Z. The RanGTP•Kap114•H2A-H2B complex used Kap114 h1-h15 and RanGTP from Kap114•RanGTP model, and Kap114 h16-h20 and H2A-H2B from Kap114•H2A-H2B. All the models were roughly docked into the map using UCSF Chimera (52) before subjected to real-space refinement with global minimization and rigid body restraints on Phenix (53). This is followed by manually rounds of rebuilding using Coot (54), and ISOLDE (55) on UCSF ChimeraX (56) and subjected to last round of refinement in Phenix. Structure interfaces were analyzed using ENDscript 2.0 (57) with contacts cut-off of 4 Å. These contacts were then manually curated and mapped onto a multiple sequence alignment generated by MAFFT (58) and visualized by ESPript 3.0 (57). We used PyMOL version 2.5 for 3D structure analysis (59).

### Nucleosome Assembly Assays

Tetrasomes containing *Xl*(H3-H4)2 and 147 bp Widom 601 DNA were reconstituted as described in Dyer et al (60). To monitor nucleosome assembly, tetrasomes were held constant at 1.25 μM and H2A-H2B or pre-formed complexes of Imp9•H2A-H2B (1:1), RanGTP•Imp9•H2A-H2B (1:1:1), Kap114•H2A-H2B (1:1), or RanGTP•Kap114•H2A-H2B (1:1:1) were titrated at 2 and 3 molar equivalents. Proteins were dialyzed into the same buffer overnight (20 mM HEPES, pH 7.5, 150 mM NaCl, 2 mM MgAcetate, 2 mM TCEP, 10% (v/v) glycerol). Samples were separated by 5% native PAGE. Gels were run for 75 min at 150 V at 4 °C in 0.5x TBE. Gels were stained with ethidium bromide and then Coomassie. Gel shown was one of ≥ 3 repeats.

## Acknowledgments

We thank Abhilash Padavannil for the important biochemical studies that led to this work. A portion of this research was supported by NIH grant U24GM129547 and performed at the PNCC at OHSU and accessed through EMSL (grid.436923.9), a DOE Office of Science User Facility sponsored by the Office of Biological and Environmental Research. We thank the Structural Biology Laboratory and the Cryo-EM Facility at UTSW, which are partially supported by grant RP170644 from the Cancer Prevention & Research Institute of Texas (CPRIT), for cryo-EM studies for their assistance with cryo-EM data collection. We thank Chad Brautigam and the Macromolecular Biophysics Resource at UTSW for training and use of their analytical ultracentrifuge. We also thank the Erzberger lab for the use of their equipment for fluorescence polarization experiments. We acknowledge The Histone Source at Colorado State University for generating the *Xl* and *Sc* histone H2A and H2B proteins used in this study. This work was funded by NIGMS of NIH under Awards R35GM141461 (Y.M.C.), R01GM069909 (Y.M.C.), R35GM133751 (S.D.), T32GM008203 (J.J.), the Welch Foundation Grants I-1532 (Y.M.C.), AT-2059-20210327 (S.D.), NSF MRI 2018188 (S.D.), support from the Alfred and Mabel Gilman Chair in Molecular Pharmacology, Eugene McDermott Scholar in Biomedical Research (Y.M.C.), Mary Kay International Postdoctoral Fellowship (N.E.B.) and the Gilman Special Opportunities Award (H.Y.J.F.).

## Supplemental Information

**Figure S1.**
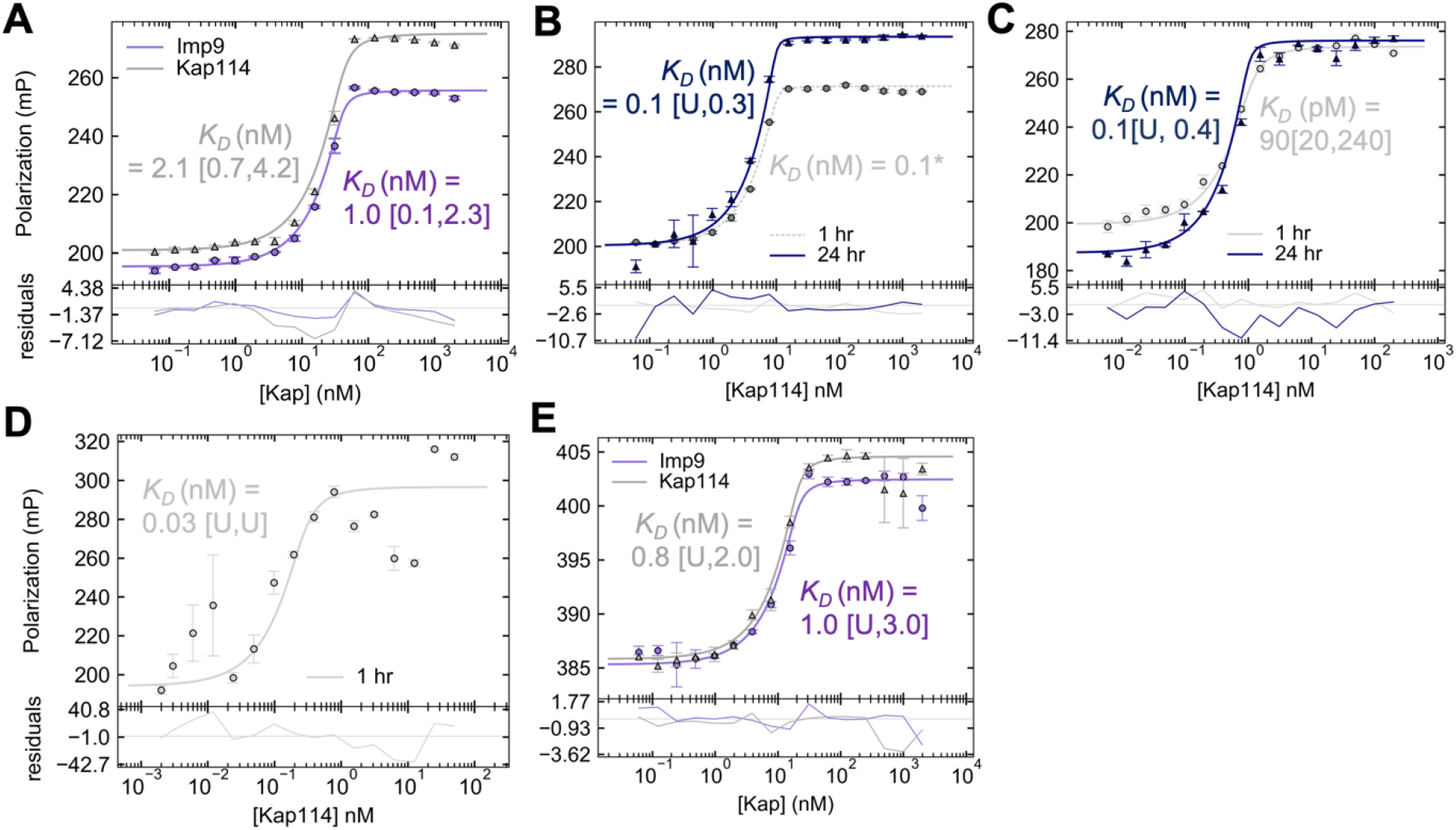
FP analysis of Kap114 and Imp9 binding to H2A-H2B or RanGTP. **(A)** Kap114 (in purple) or Imp9 (in gray) titrated into 40 nM ^XFD488^H2A-H2B. Data points represent triplicate measurements and standard deviation as error bars. Data fitted with 1-site binding model is shown in solid line. Binding affinity is indicated on the graph with 95% confidence interval shown in brackets. The curve fit is not optimal, and the binding is too tight for K_D_ obtained to be accurate with the concentration of H2A-H2B used, thus probe concentration was lowered to **(B)** 10 nM, **(C)** 1 nM and **(D)**0.25nM ^XFD488^H2A-H2B. Measurement was taken at 1 hr (gray) and 24 hr later (marine). For the 1 hr measurement of **(B)**, data could not be fitted, and the dotted line represents predicted data with a K_D_ of 0.1 nM. Gain was readjusted to accommodate for lower concentration of fluorophore used. From the noise level in **(D)**, it can be seen that the gain here is too high and therefore 1 nM in **(C)** is likely the lower limit for this plate reader. Therefore, we could not accurately measure the binding affinity as the ^XFD488^H2A-H2B used is still higher than that of the K_D_. However, it is likely that the K_D_ is < 0.1 nM. **(E)** Kap114 (in purple) or Imp9 (in gray) titrated into 20 nM mNeoGreen-RanGTP.

**Figure S2.**
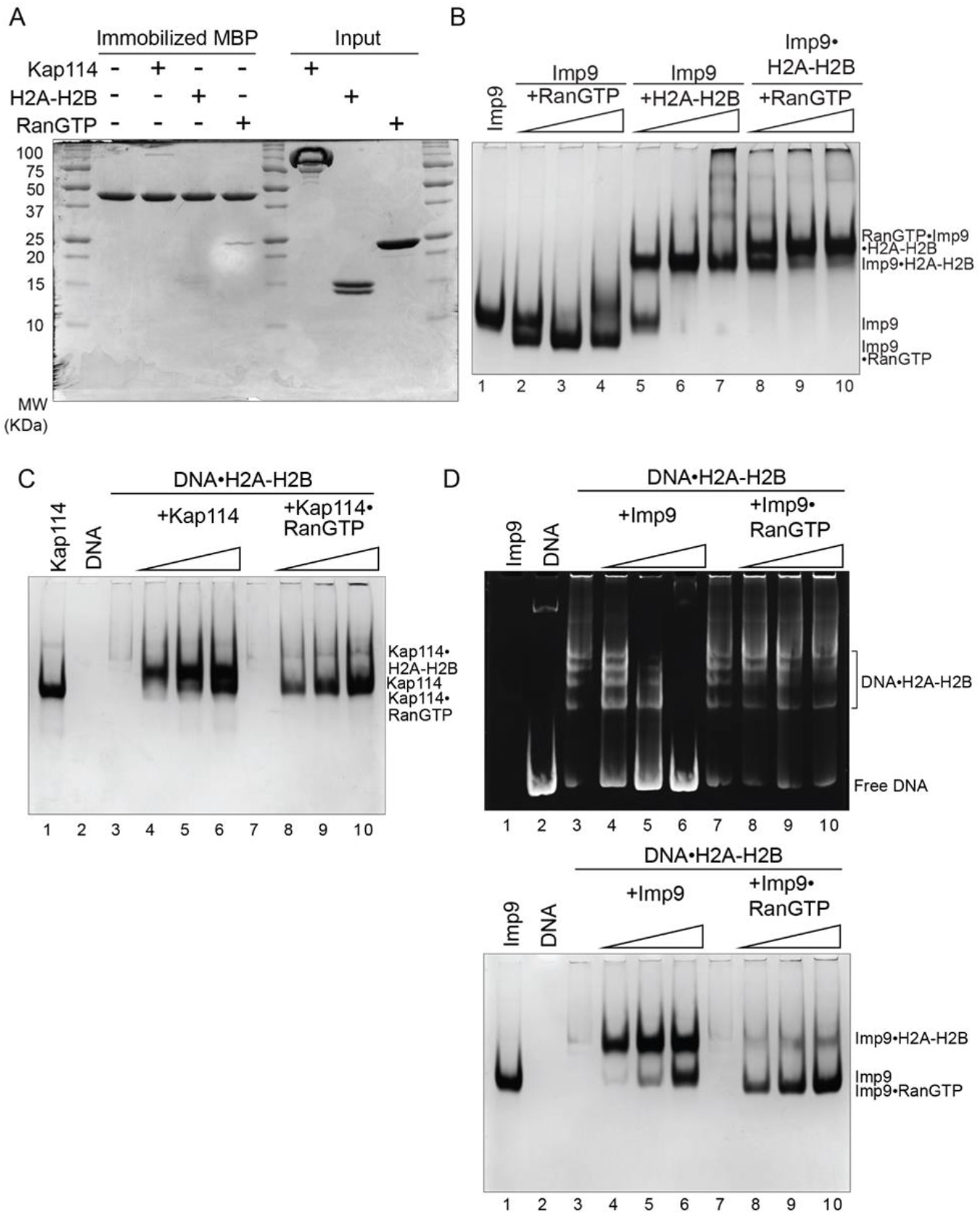
Kap114 and Imp9 binding to H2A-H2B and RanGTP and DNA competition assays. (**A**) Pull-down binding assays of immobilized MBP on amylose beads with untagged Kap114, H2A-H2B, and RanGTP added. After extensive washing, bound and input proteins were visualized by Coomassie-stained SDS-PAGE. (**B**) Constant Imp9 (lanes 2-7) was titrated with 0.5, 1, or 1.5 molar ratio of RanGTP (lanes 2-4) or H2A-H2B (lane 5-7). Constant Imp9•H2A-H2B was titrated with 0.5, 1, or 1.5 molar ratio of RanGTP (lanes 8-10). Protein was visualized by Coomassie-stained native PAGE. (**C**) DNA competition assay with Kap114 (lanes 4-6) titrated at 0.5, 1, or 1.5 molar equivalents of H2A-H2B (in a DNA•H2A-H2B 1:7 complex), while Kap114•RanGTP (1:1, lanes 8-10) is titrated at 0.25, 0.5, or 1.0 molar equivalents of H2A-H2B (in a DNA•H2A-H2B 1:7 complex. Native PAGE was visualized with Coomassie staining. Ethidium bromide is shown in Figure 1D. (**D**) DNA competition assay with Imp9 (lanes 4-6) titrated at 0.5, 1, or 1.5 molar equivalents of H2A-H2B (in a DNA•H2A-H2B 1:7 complex), while Imp9•RanGTP (1:1, lanes 8-10) is titrated at 0.25, 0.5, or 1.0 molar equivalents of H2A-H2B (in a DNA•H2A-H2B 1:7 complex. Native PAGE was visualized with ethidium bromide (top) and Coomassie staining (bottom).

**Figure S3.**
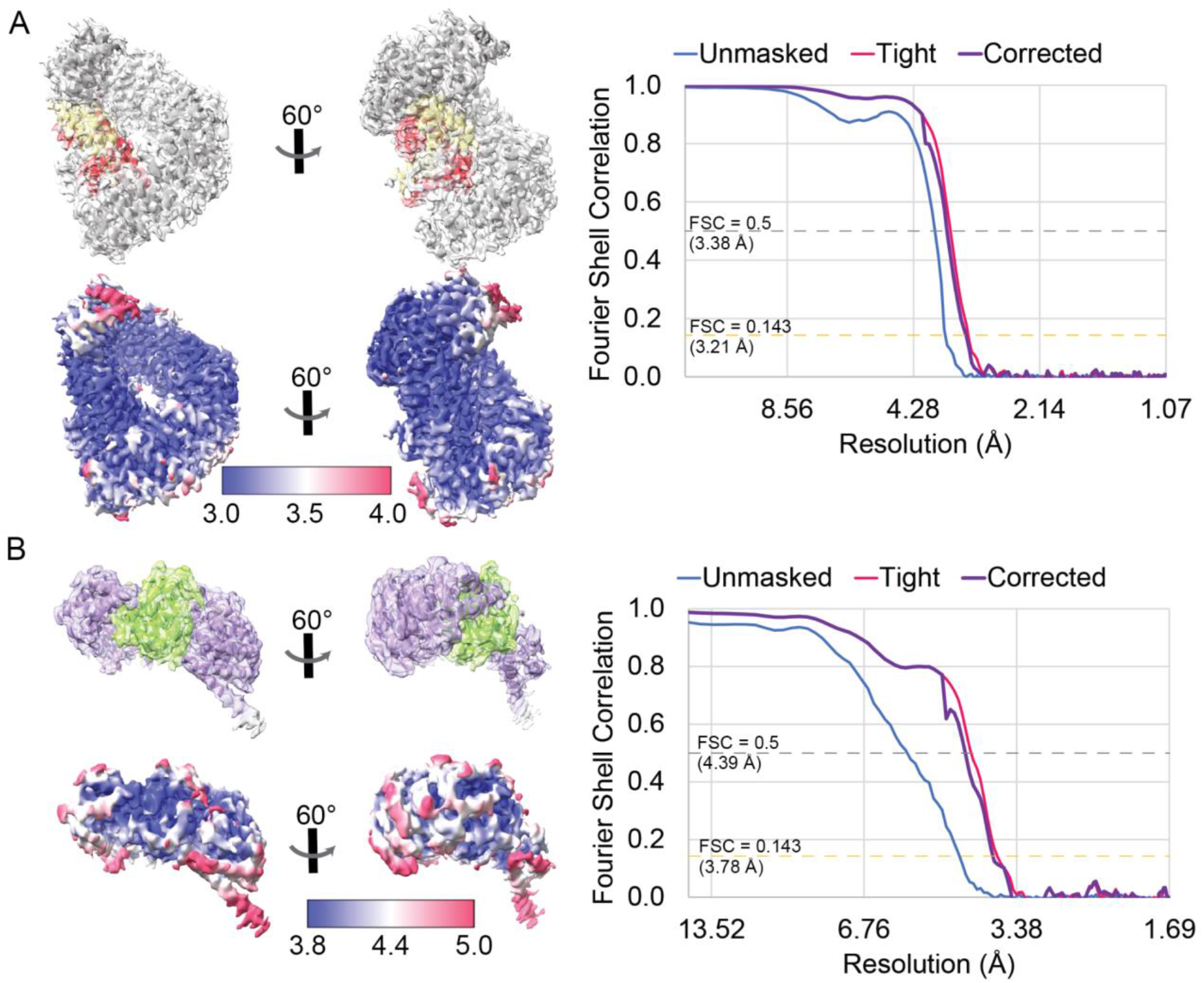
Cryo-EM map statistics of Kap114•H2A-H2B and Imp9•RanGTP. (**A**) The Kap114•H2A-H2B model, colored gray•yellow-red, is docked into its map with the local resolution displayed below (left). The local resolution was colored to show according to the scale displayed. The FSC curves were plotted with unmasked in blue, tight mask in pink, and corrected mask in purple (right). The gold dashed line marks the FSC = 0.143, and the gray dashed line represents FSC = 0.5. (**B**) Imp9•RanGTP model, colored purple•green, is docked into its map - shown with its corresponding local resolution and its FSC curve.

**Figure S4.**
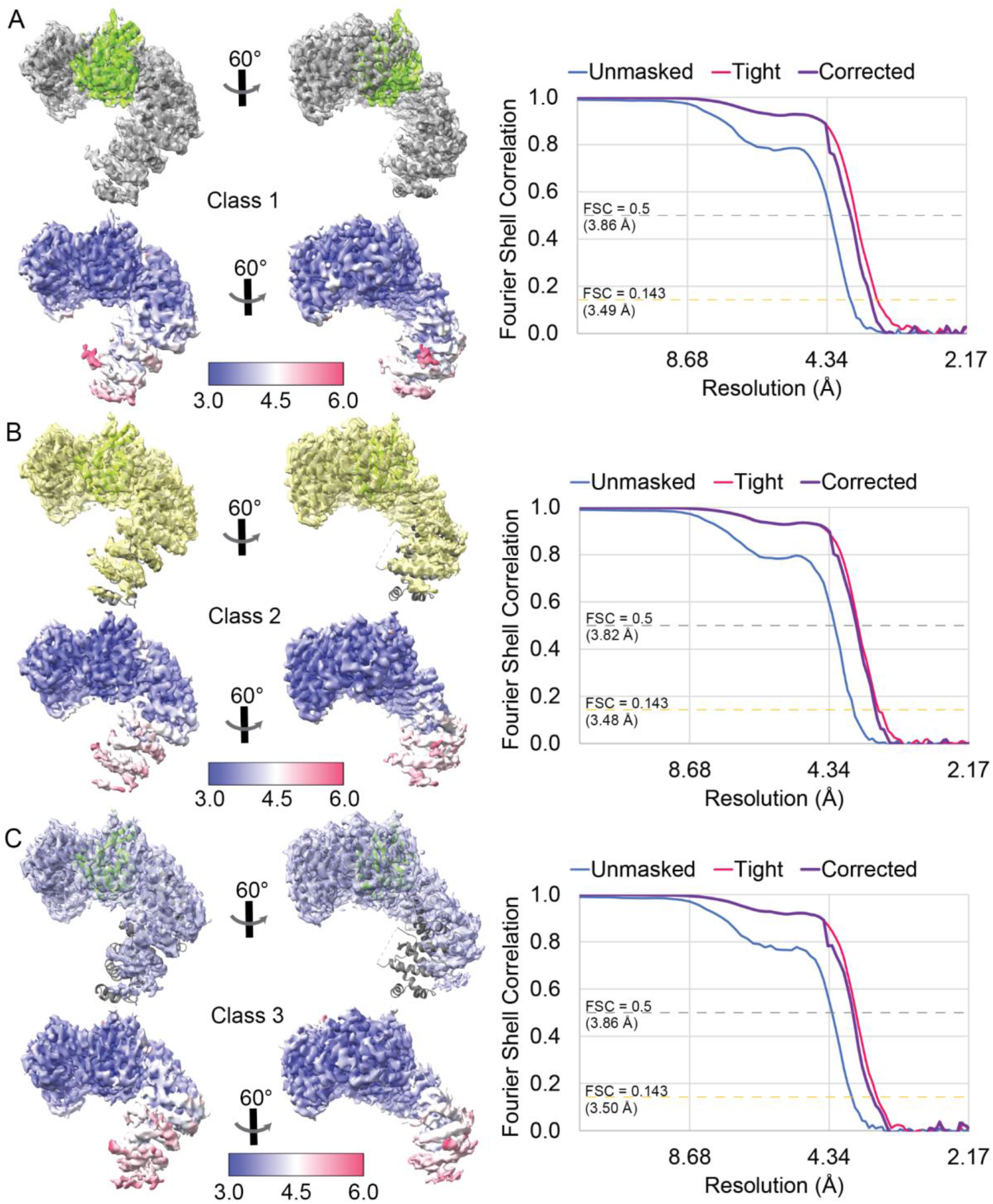
Cryo-EM map statistics of additional classes for Kap114•RanGTP. (**A**) The Kap114•RanGTP model of class 1, colored gray•green, is docked into the class 1 map with the local resolution displayed below (left). The local resolution was colored according to the scale displayed. The FSC curves were plotted with unmasked in blue, tight mask in pink, and corrected mask in purple (Right). The gold dashed line marks the FSC = 0.143, and the gray dashed line represents FSC = 0.5. The same class 1 model is docked into (**B**) a yellow class 2 map and (**C**) a light blue class 3 map - shown with the corresponding local resolution and FSC curve.

**Figure S5.**
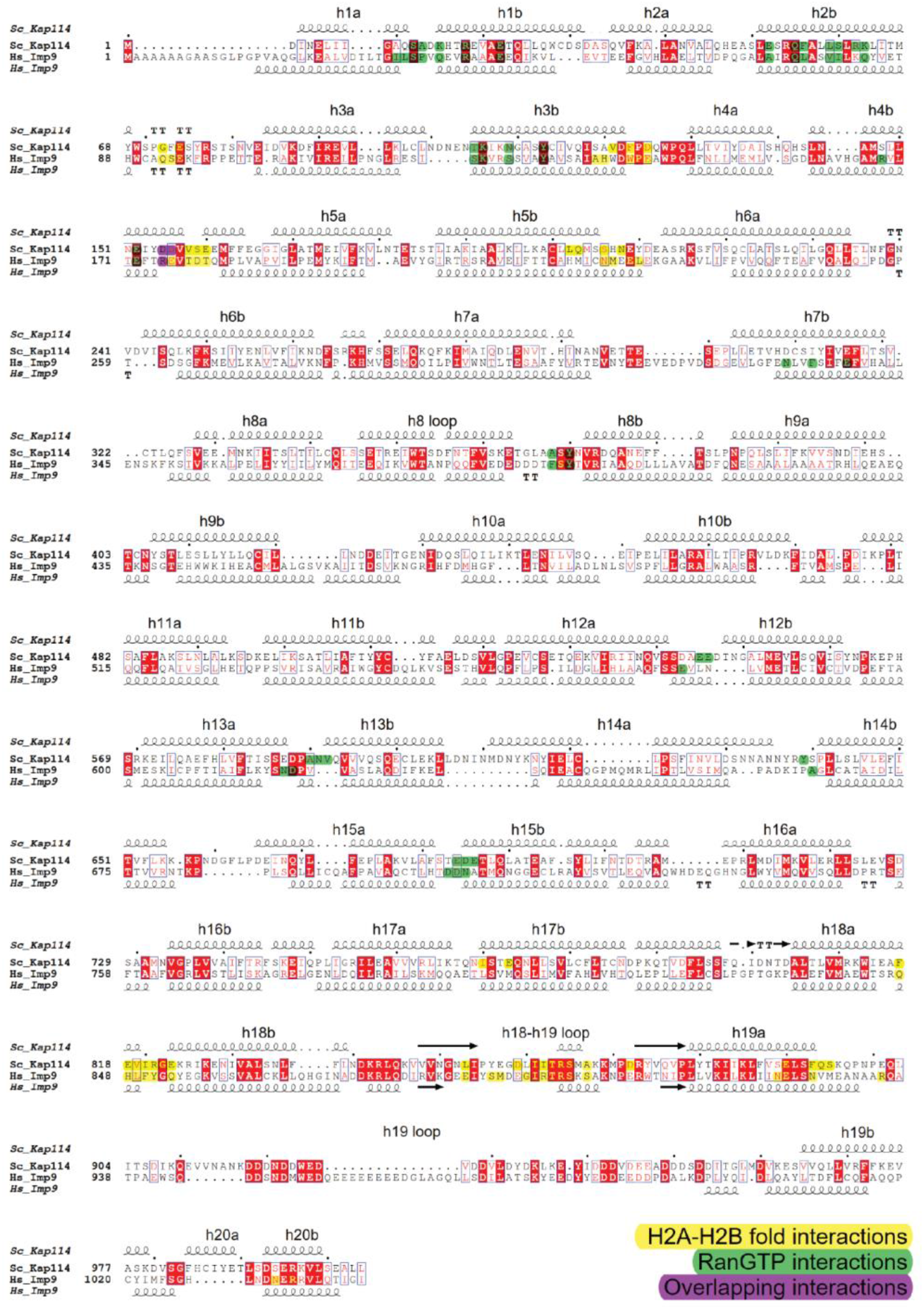
Sequence alignment of Kap114 and Imp9 generated using MAFFT. ESPript 3.0 was used to visualize the alignment with the secondary structures annotated from the histone-bound binary structures. The boxed residues in red represent identical residues, and the ones outlined in blue represent similar residues. The HEAT repeats are labeled above Kap114 secondary structure, and contacts in the binary structures are highlighted with H2A-H2B core interactions in yellow, RanGTP interactions in green, and the overlapping interactions by both binary structures in purple.

**Figure S6.**
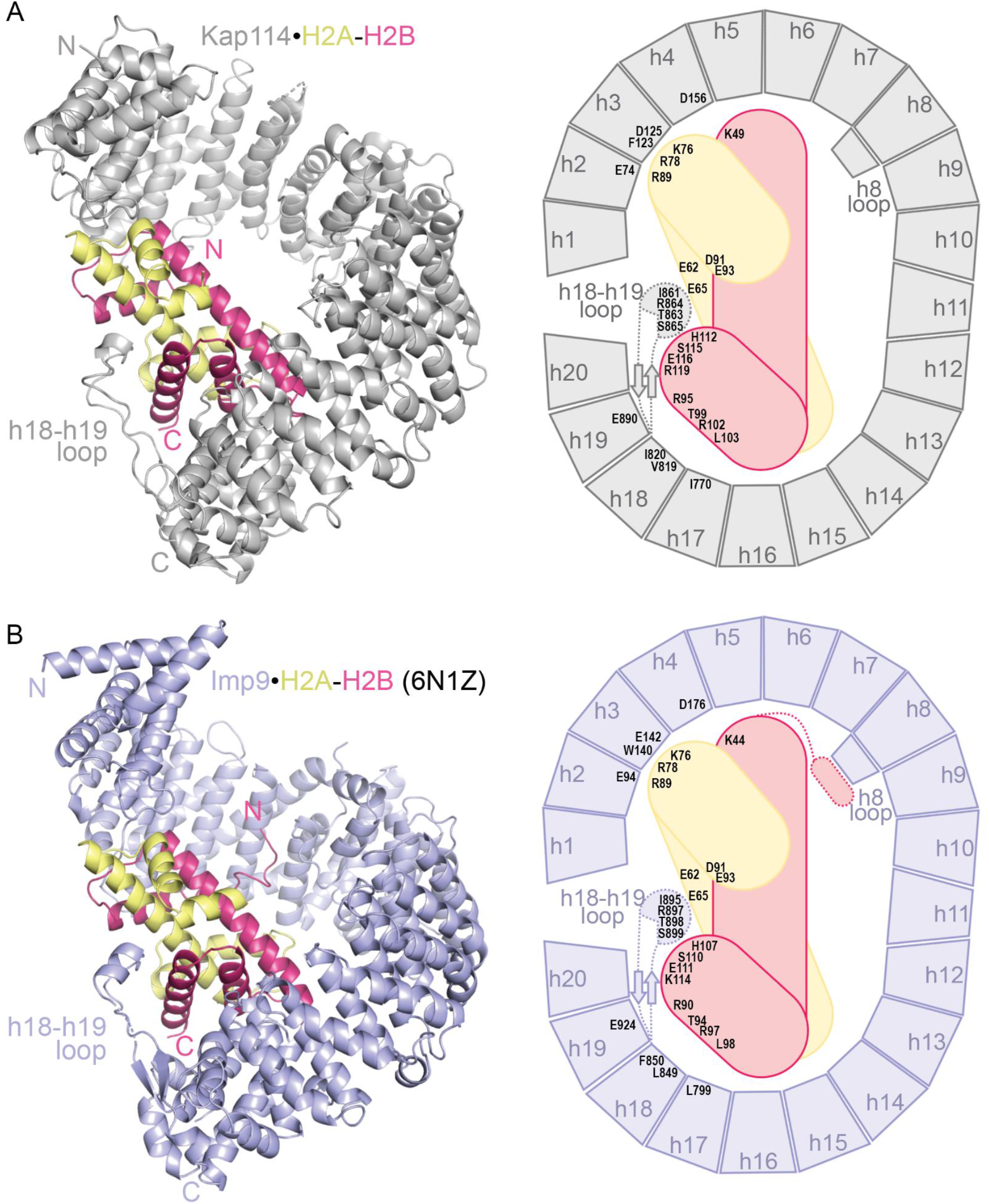
Conservation of Kap114 and Imp9 binding H2A-H2B. (**A**) A cartoon of Kap114•H2A-H2B colored in gray•yellow-red (left). A schematic with the same color scheme depicts both topology and the conserved residues Imp9 and Kap114 share when contacting H2A-H2B (right). (**B**) A cartoon of Imp9•H2A-H2B colored in light purple•yellow-red and its schematic.

**Figure S7.**
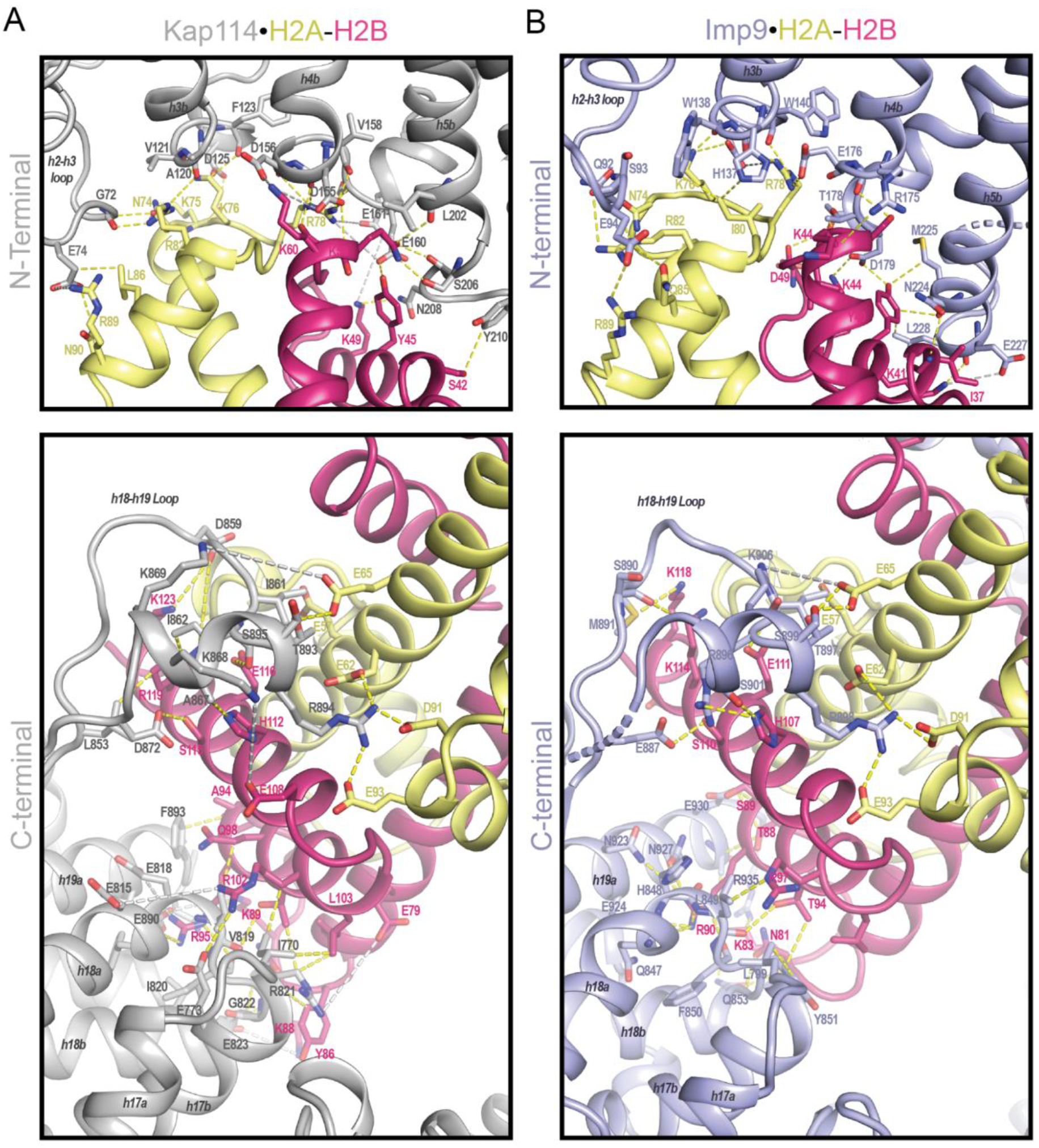
Kap114 and Imp9 interactions with H2A-H2B. **(A)** Detailed interactions of Kap114•H2A-H2B and **(B)** Imp9•H2A-H2B complex at the N- and C-terminal interface of the importins with H2A-H2B. In all panels, Kap114 is gray, Imp9 is light purple, H2A is yellow, and H2B is red. The yellow and gray dotted lines represent interactions under <4 Å (yellow) and long-range electrostatic interactions under <8 Å (gray), respectively.

**Figure S8.**
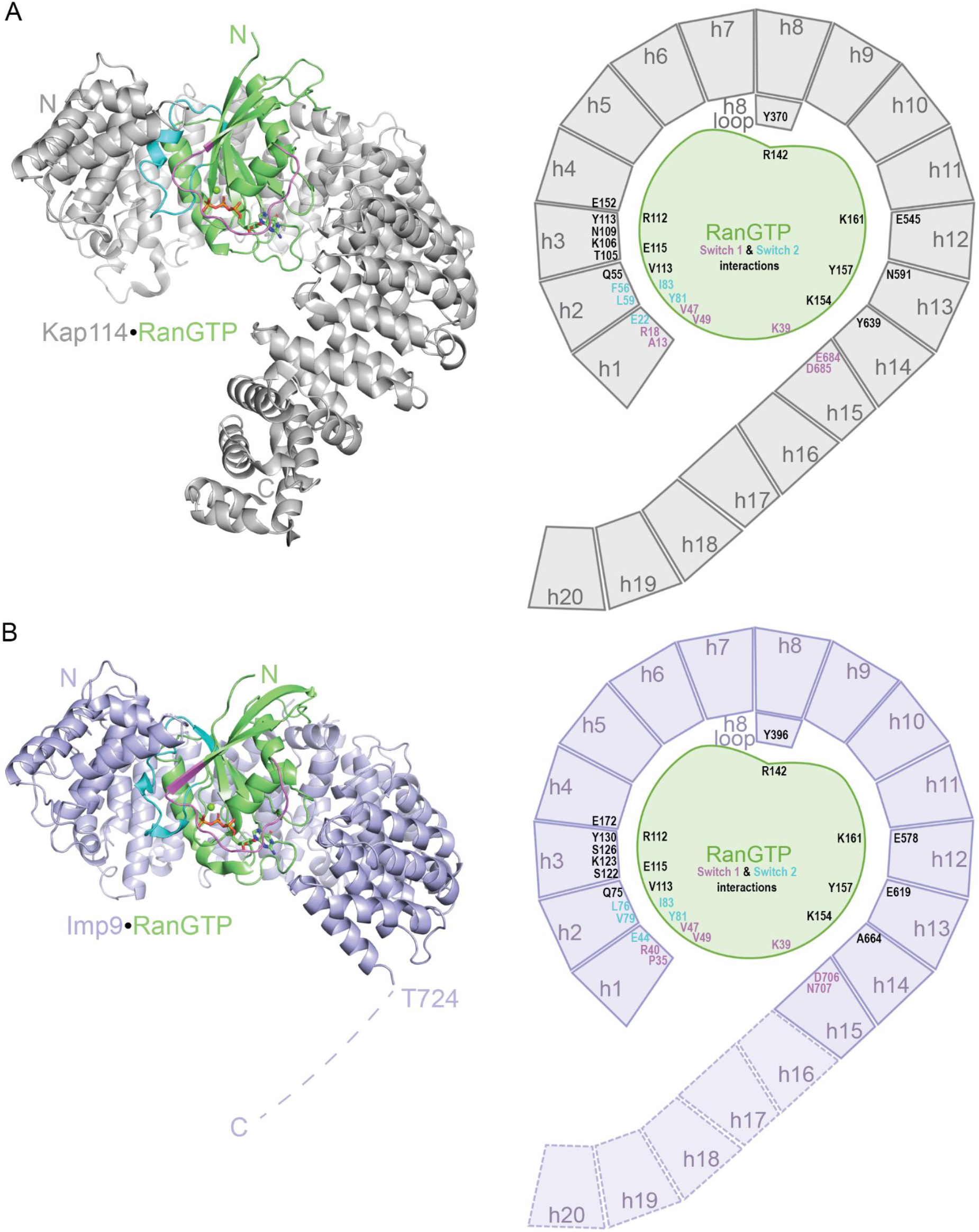
Conservation of Kap114 and Imp9 binding RanGTP. (**A**) A cartoon of Kap114•RanGTP colored in gray•green (Left). A schematic with the same color scheme depicts both topology and the conserved residues Imp9 and Kap114 share when contacting RanGTP (Right). (**B**) A cartoon of Imp9•RanGTP colored in light purple•green and its schematic.

**Figure S9.**
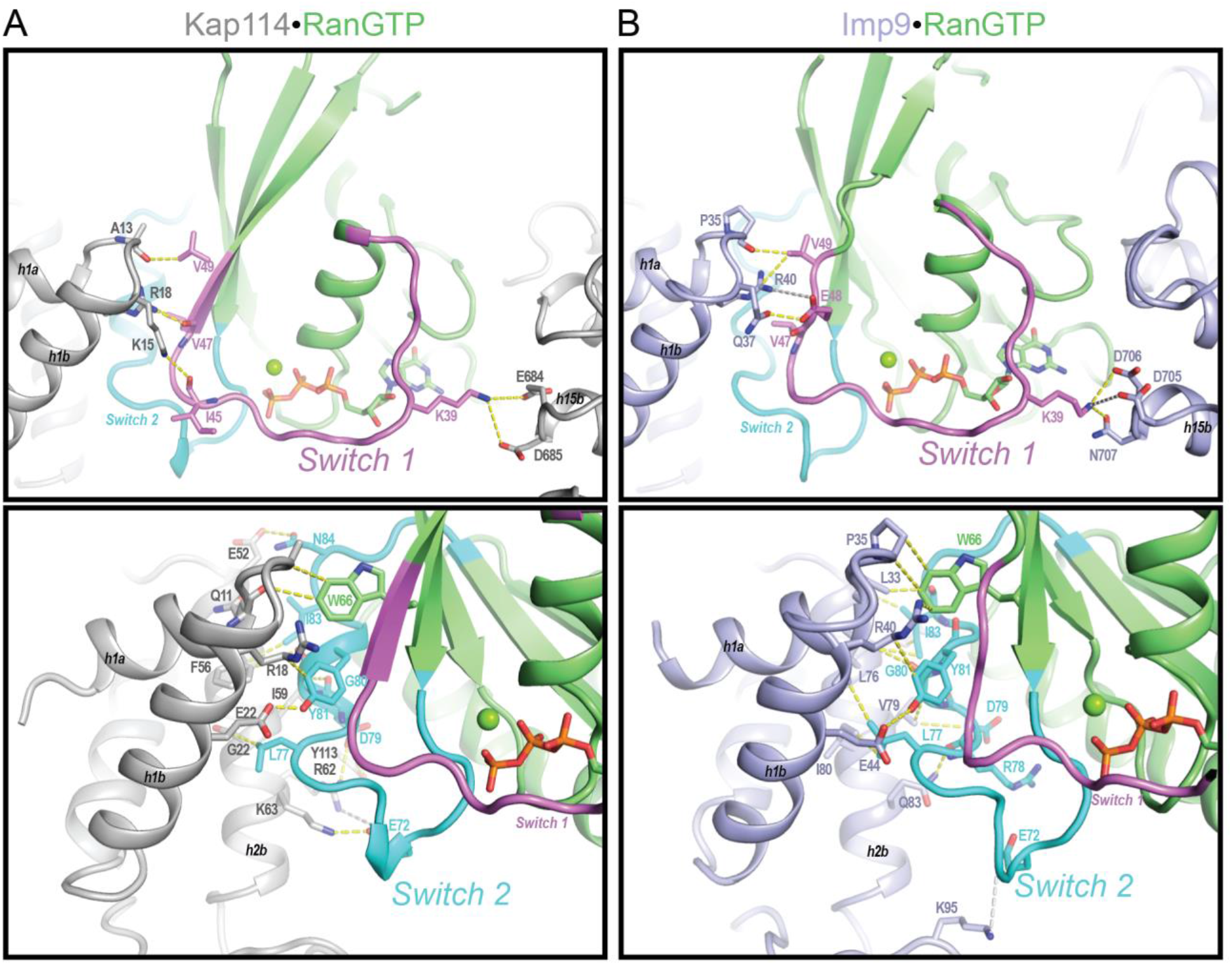
Kap114 and Imp9 interactions with RanGTP. **(A)** Detailed interactions of Kap114•RanGTP and **(B)** Imp9•RanGTP complex at the Switch 1 and Switch 2 interface of the RanGTP with Kaps. In all panels, Kap114 is gray, Imp9 is light purple, RanGTP is in green with switch 1 colored violet and switch two colored cyan. The yellow and gray dotted lines represent interactions under <4 Å (yellow) and long-range electrostatic interactions under <8 Å (gray), respectively.

**Figure S10.**
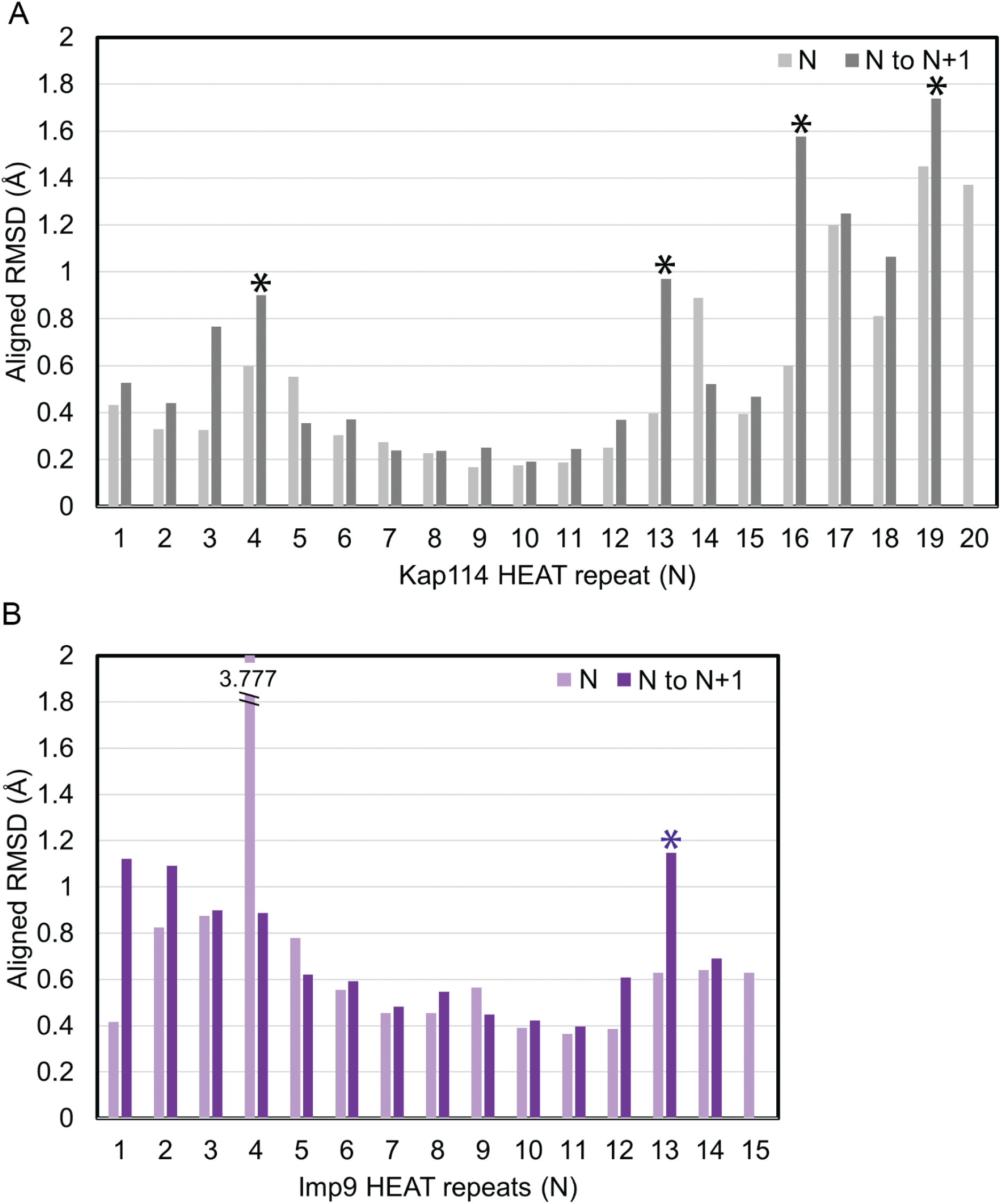
Alignment of Kap114 or Imp9 bound to RanGTP versus H2A-H2B. **(A)** The HEAT repeats of the binary Kap114 complexes and **(B)** binary Imp9 complexes were aligned by individual HEAT repeats, N, and paired HEAT repeats, N to N+1. The asterisk denotes local maximum RMSD between the paired HEAT repeats that corresponds to a hinge.

**Figure S11.**
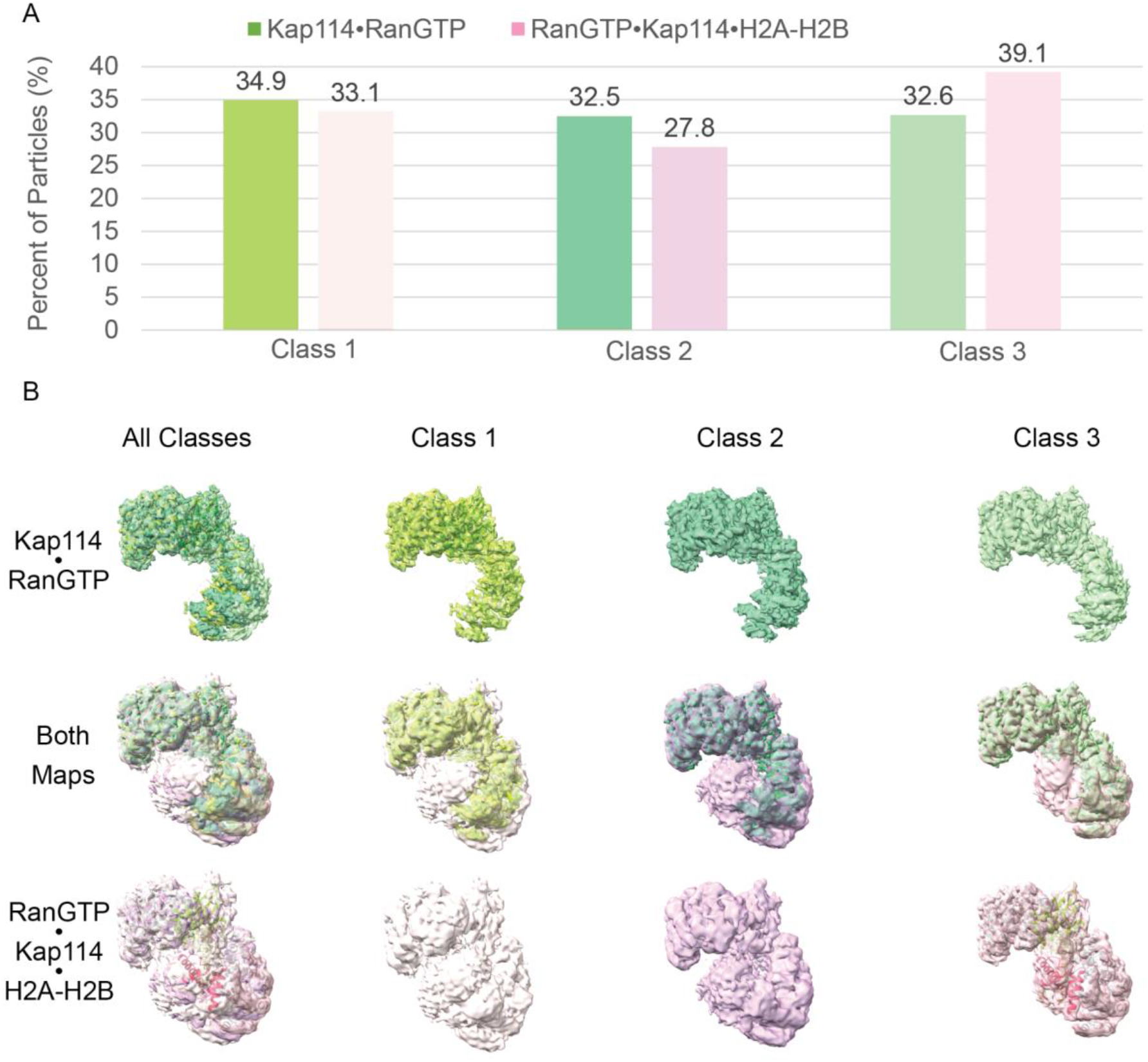
Populations of three distinct classes of Kap114•RanGTP and RanGTP•Kap114•H2A-H2B. **(A)** The percentage of particles in each class are plotted in a bar chart with Kap114•RanGTP in green and RanGTP•Kap114•H2A-H2B in pink. **(B)** A visual overlay of the maps obtained from the three classes for the binary complex in green and the ternary complex in pink. Where class1 exhibits the smallest degree of opening, class 3 the widest opening of the three classes, and class 2 the moderate opening of the Kap114 C-terminal repeats.

**Figure S12.**
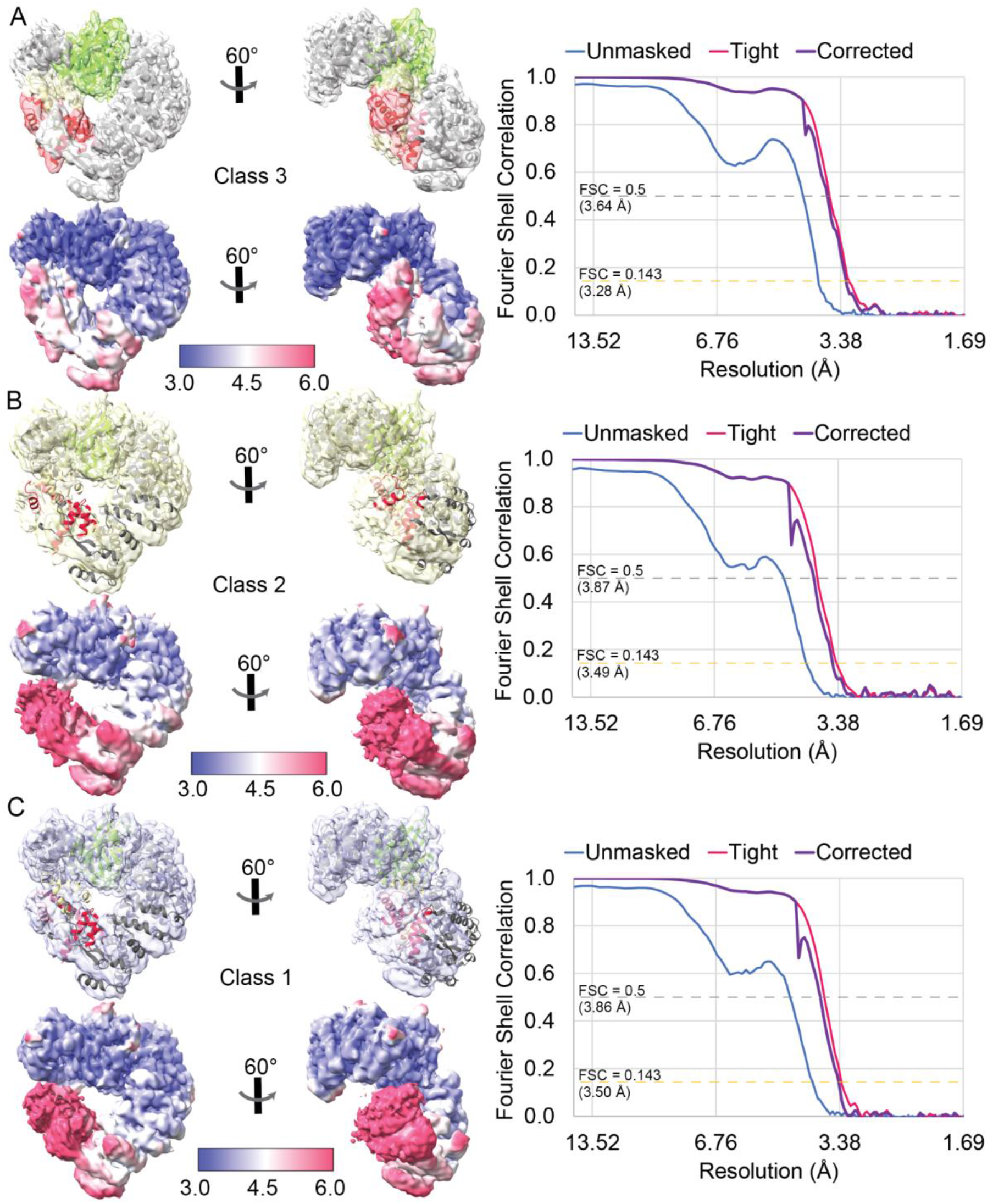
Cryo-EM map statistics of additional classes for RanGTP•Kap114•H2A-H2B. (**A**) The RanGTP•Kap114•H2A-H2B model of class 3, colored green•gray•yellow-red, is docked into the class 3 map with the local resolution displayed below (left). The local resolution was colored according to the scale displayed. The FSC curves were plotted with unmasked in blue, tight mask in pink, and corrected mask in purple (right). The gold dashed line marks the FSC = 0.143, and the gray dashed line represents FSC = 0.5. The same class 3 model is docked into (**B**) a yellow class 2 map and (**C**) a light blue class 1 map - shown with the corresponding local resolution and FSC curve.

**Figure S13.**
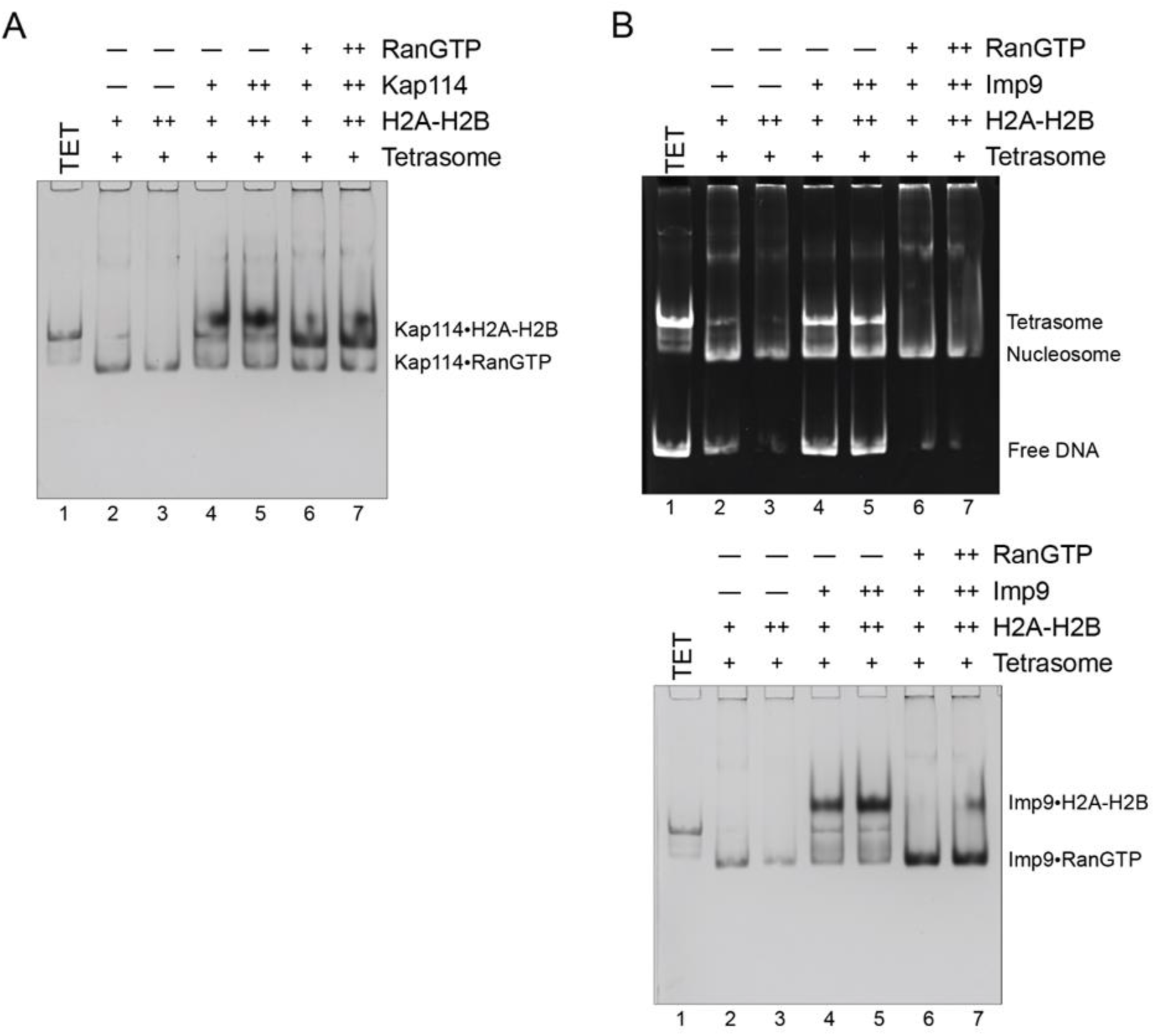
Nucleosome assembly assays. (**A**) Nucleosome assembly assay where either H2A-H2B, Kap114•H2A-H2B or RanGTP•Kap114•H2A-H2B is titrated in molar equivalents of 2.0 and 3.0 to tetrasome (TET; 1.25 μM). Native page gel was Coomassie-stained. Ethidium bromide-stained gel is in *Figure 4A*. (**B**) Nucleosome assembly assay where either H2A-H2B, Imp9•H2A-H2B or RanGTP•Imp9•H2A-H2B is titrated in molar equivalents of 2.0 and 3.0 to tetrasome (TET; 1.25 μM). Images of the same native gel, ethidium bromide-stained (top) and Coomassie-stained (bottom) are aligned for comparison.

**Table S1.** Kap114 Cryo-EM data collection, refinement, and validation statistics

|  | Kap114·H2A-H2B<br>EMD-28788<br>PDB 8F0X | RanGTP·Kap114<br>EMD-28782<br>PDB 8F19 | RanGTP·Kap114·H2A-H2B<br>EMD-28796<br>PDB 8F1E |
| --- | --- | --- | --- |
| **Data collection** |  |  |  |
| Facility | PNCC | PNCC | UTSW |
| Magnification | 81kx | 81kx | 105kx |
| Voltage (kV) | 300 | 300 | 300 |
| Electron exposure (e<sup>-</sup>/Å<sup>2</sup>) | 50 | 50 | 52 |
| Defocus range (μm) | 1.5-2.5 | 1.5-2.5 | 1.5-2.5 |
| Pixel size (Å) | 0.5295 | 0.5295 | 0.415 |
| Symmetry imposed | C1 | C1 | C1 |
| Initial particle images (no.) | 6,632,116 | 7,046,263 | 3,401,608 |
| Final particle images (no.) | 554,529 | 277,305 | 134,915 |
| Map resolution (Å) | 3.21 | 3.49 | 3.28 |
| FSC threshold | 0.143 | 0.143 | 0.143 |
| **Refinement** |  |  |  |
| Initial model used (PDB) | 6AHO, 6N1Z | 6AHO, 3W3Z | 8F0X, 8F19 |
| Model resolution (Å) | 2.5/3.0/3.3 | 2.8/3.2/3.6 | 3.1/3.2/3.5 |
| FSC threshold | 0/0.143/0.5 | 0/0.143/0.5 | 0/0.143/0.5 |
| Map sharpening *B* factor | 158.4 | 165.0 | 126.5 |
| Model composition |  |  |  |
| Nonhydrogen atom | 8,743 | 8,717 | 10,174 |
| Protein residues | 1,101 | 1,087 | 1,273 |
| Ligand | --- | 2 (GTP; Mg) | 2 (GTP; Mg) |
| *B* factors (Å<sup>2</sup>) |  |  |  |
| Protein | 58.65 | 70.40 | 69.99 |
| Ligand | --- | 60.23 | 138.41 |
| R.m.s. deviations |  |  |  |
| Bond lengths (Å) | 0.004 | 0.003 | 0.003 |
| Bond angles (°) | 0.647 | 0.667 | 0.668 |
| **Validation** |  |  |  |
| MolProbity score | 1.40 | 1.43 | 1.58 |
| Clashscore | 7.37 | 7.81 | 11.76 |
| Poor rotamers (%) | 0 | 0 | 0.17 |
| Ramachandran plot |  |  |  |
| Favored (%) | 98.36 | 99.07 | 98.26 |
| Allowed (%) | 1.74 | 0.93 | 1.74 |
| Disallowed (%) | 0 | 0 | 0 |

**Table S2.** Imp9 Cryo-EM data collection, refinement, and validation statistics

|  | Imp9-RanGTP<br>EMD-28899<br>PDB 8F7A |
| --- | --- |
| **Data collection** |  |
| Facility | UTSW |
| Magnification | 105kx |
| Voltage (kV) | 300 |
| Electron exposure (e<sup>-</sup>/Å<sup>2</sup>) | 52 |
| Defocus range (μm) | 1.5-2.5 |
| Pixel size (Å) | 0.417 |
| Symmetry imposed | C1 |
| Initial particle images (no.) | 2,198,920 |
| Final particle images (no.) | 239,496 |
| Map resolution (Å) | 3.78 |
| FSC threshold | 0.143 |
| **Refinement** |  |
| Initial model used (PDB) | 6N1Z, 3W3Z |
| Model resolution (Å) | 3.3/3.5/4.0 |
| FSC threshold | 0/0.143/0.5 |
| Map sharpening *B* factor (Å<sup>2</sup>) | 204.4 |
| Model composition |  |
| Nonhydrogen atom | 6,721 |
| Protein residues | 849 |
| Ligand | 2 (GTP; Mg) |
| *B* factors (Å<sup>2</sup>) |  |
| Protein | 59.02 |
| Ligand | 80.04 |
| R.m.s. deviations |  |
| Bond lengths (Å) | 0.003 |
| Bond angles (°) | 0.611 |
| **Validation** |  |
| MolProbity score | 1.80 |
| Clashscore | 10.24 |
| Poor rotamers (%) | 0 |
| Ramachandran plot |  |
| Favored (%) | 96.06 |
| Allowed (%) | 3.82 |
| Disallowed (%) | 0.12 |

## Notes

**Competing Interest Statement:** No competition interest declared.

### Competing Interest Statement

The authors have declared no competing interest.

### Summary of Updates

517512 was divided into two separate non-overlapping but complementary manuscripts.

